# Circulating metabolites and the risk of type 2 diabetes: a prospective study of 11,896 young adults from four Finnish cohorts

**DOI:** 10.1101/513648

**Authors:** Ari V. Ahola-Olli, Linda Mustelin, Maria Kalimeri, Johannes Kettunen, Jari Jokelainen, Juha Auvinen, Katri Puukka, Aki S. Havulinna, Terho Lehtimäki, Mika Kähönen, Markus Juonala, Sirkka Keinänen-Kiukaanniemi, Veikko Salomaa, Markus Perola, Marjo-Riitta Järvelin, Mika Ala-Korpela, Olli Raitakari, Peter Würtz

## Abstract

**Objective:** Advances in metabolomics now allow high-throughput biomarker profiling of large population studies. We aimed to identify circulating metabolic biomarkers predictive of type 2 diabetes in young adults.

**Methods:** Nuclear magnetic resonance metabolomics was used to quantify 229 metabolic measures in 11,896 individuals from four Finnish cohorts (mean age 33 years, range 24–45). Associations between baseline metabolites and risk of type 2 diabetes onset during 8–15 years of follow-up (392 incident cases) were assessed by logistic regression adjusted for sex, age, body mass index, and fasting glucose.

**Results:** Out of 229 metabolic measures, 113 were associated with incident diabetes in meta-analysis of the four cohorts (P<0.0009; odds ratios per 1-SD: 0.59–1.50). Among the strongest predictors of diabetes risk were branched-chained and aromatic amino acids (odds ratios 1.31–1.33), triglyceride fractions within the largest very-low-density lipoprotein particles (VLDL; odds ratios 1.33–1.50)), as well as linoleic omega-6 fatty acids (odds ratio 0.75) and free cholesterol in large high-density lipoprotein particles (HDL; odds ratio 0.59). A biomarker score comprised of phenylalanine, free cholesterol in large HDL, and the ratio of cholesteryl esters to total lipids in large VLDL was predictive of the risk for future diabetes in an independent validation cohort (odds ratio 10.1 [95% confidence intervals 4.2-24.1] comparing individuals in upper *vs* lower fifth of biomarker score). Adjustment for routine lipids and insulin attenuated the odds ratio to 5.8 [2.2-15.1].

**Conclusions:** Metabolic aberrations across multiple molecular pathways are predictive of the long-term risk of type 2 diabetes in young adults. Comprehensive metabolic profiling may potentially help targeting preventive interventions for young asymptomatic individuals at increased risk for type 2 diabetes.

## INTRODUCTION

The global prevalence of type 2 diabetes is increasing rapidly, particularly in low-income and middle-income countries (1). Type 2 diabetes is associated with increased mortality from vascular and numerous other causes, and reduced quality of life, causing an immense societal cost burden (2, 3). Given the availability of lifestyle interventions that are effective at preventing or delaying the onset of type 2 diabetes (4, 5), early identification of individuals at high risk is important. The risk for developing type 2 diabetes is to some extent reflected in current laboratory measures, including hyperglycaemia and dyslipidemia. Such metabolic aberrations are often present years before the diagnosis can be set. This has led to widespread implementation of prediabetes screening programs for identifying those at high risk for progression to diabetes. However, current screening approaches are ineffective in identifying high risk individuals, leading to missed opportunities for targeted prevention (6).

Detailed metabolic profiling technologies, also known as metabolomics, are increasingly used in epidemiological studies (7, 8). This has proven a powerful approach to identify biochemical changes occurring before disease onset and elucidate the pathophysiology of type 2 diabetes. Multiple nested case-control studies have identified circulating metabolites associated with the risk for type 2 diabetes using a range of technological assays, based on mass spectrometry or nuclear magnetic resonance (NMR) (7, 9, 10). The most consistent biomarker associations with type 2 diabetes have been observed for branched-chain and aromatic amino acids (8). Genetic evidence and experimental data suggest that these amino acids may be causally implicated in the development of insulin resistance and type 2 diabetes (11, 12). However, previous metabolomics studies have commonly involved a modest number of participants, and have almost exclusively been conducted for middle-aged and older individuals.

In this study, we characterized a comprehensive metabolic signature of increased risk for future type 2 diabetes in young adults, up to 15 years before disease onset. We used NMR metabolomics to quantify 229 metabolic measures in 11,896 individuals from four population-based cohorts with individuals aged 24–45 at blood draw. We aimed to describe the metabolic aberrations reflective of the risk for future type 2 diabetes in young adults, including their consistency between men and women and across cohorts. In addition, we explored the predictive value of a multi-biomarker score for early risk of type 2 diabetes.

## METHODS

### Study populations

The study involved 11,896 individuals from four prospective population-based cohorts in Finland. An overview of the study cohorts and participants included in the present analyses is shown in **Supplementary Figure 1**. Details of the individual cohorts are provided in **Supplementary Methods**. In all cohorts, we excluded individuals with diabetes at baseline, pregnant women, study participants aged over 45 years at the blood draw, and those lacking follow-up information on diabetes diagnosis. Briefly, the characteristics of each cohort were as follows:

1. In the Cardiovascular Risk in <u>Y</u>oung <u>F</u>inns <u>S</u>tudy (YFS), serum metabolites were quantified from 2,248 individuals in the 2001 survey. The final sample consisted of 2,141 individuals of age range 24–39 years. Follow-up time was 10 years. Type 2 diabetes diagnoses at 10-year follow-up were based either on HbA1c or fasting glucose assessed in the 2011 re-survey, or nationwide registers on reimbursement for diabetes medication or in-patient hospital ICD-10 diagnosis of diabetes (**Supplementary Methods**) (13).
2. In FINRISK-1997, serum metabolites were quantified from 7,603 individuals. The final sample consisted of 3,063 individuals when limiting analyses to participants aged 24–45. Follow-up time was 15 years. Type 2 diabetes diagnoses at follow-up were based on nationwide register data (14).
3. In the <u>D</u>ietary, <u>L</u>ifestyle, and <u>G</u>enetic Determinants of <u>O</u>besity and <u>M</u>etabolic Syndrome Study (DILGOM), serum metabolites were quantified from 4,816 individuals in 2007. The final sample consisted of 1,421 individuals when limiting analyses to participants in the age range 25-45. Follow-up time was 7.8 years. Type 2 diabetes diagnoses at follow-up were based on fasting glucose at the re-survey conducted in 2014 or nationwide register data.
4. In the <u>N</u>orthern <u>F</u>inland <u>B</u>irth <u>C</u>ohort (NFBC) of 1966, serum metabolites were quantified from 5,680 individuals in the 1997 survey. The final sample consisted of 5,275 individuals aged 30-32 years. Follow-up time was 15 years. Type 2 diabetes diagnoses were based on either fasting or 2-hour glucose at the 46-year follow-up conducted in 2012, or nationwide register data.

### Metabolite quantification

A high-throughput NMR metabolomics platform (Nightingale Health Ltd, Helsinki, Finland) was used to quantify 229 metabolic measures from baseline serum samples (15). This metabolite panel captures multiple metabolic pathways, including amino acids, glycolysis-related metabolites, fatty acids and detailed lipoprotein lipid profiles, covering triglycerides, total cholesterol, free cholesterol, esterified cholesterol, and phospholipids within 14 subclasses. Details of the NMR metabolomics experimentation have been described previously (15) and epidemiological applications have recently been reviewed (7).

### Statistical analyses

Due to skewness of the metabolite distributions, all metabolite concentrations were log_*e*_(metabolite+1) transformed prior to analyses. Although 229 metabolic measures in total were analyzed, the number of independent tests performed is lower due to the correlated nature of the measures (7). We calculated that 54 principal components explained 99% of the variation in the metabolic measures. Hence, we inferred statistical significance at meta-analysis P-value < 0.0009 (0.05/54). Odds ratios of 229 circulating metabolic measures with incidence of type 2 diabetes were assessed using logistic regression. Each metabolite was analyzed as predictor of diabetes in a separate model, adjusted for sex, baseline age, fasting glucose, and body mass index (BMI). We also assessed the influence of additional adjustment for fasting insulin. To facilitate comparison of the biomarker association magnitudes for measures with units and different concentration range, the odds ratios are scaled to 1-standard deviation (SD) increment in log-transformed metabolite concentration. Results from individual cohorts were combined using inverse variance-weighted fixed-effect meta-analysis. Analyses were also conducted separately for men and women.

To compare the pattern of metabolite associations for future diabetes risk with those of established risk factors, we subsequently assessed the cross-sectional associations of all metabolites with BMI, homeostatic model assessment insulin resistance (HOMA-IR) index, and fasting glucose in linear regression models adjusted for age and sex.

Lastly, we examined the predictive value of a combination of metabolites in the NFBC validation cohort based on a biomarker summary score derived in the three other cohorts (YFS, FINRISK-1997, and DILGOM; constituting approximately half of the incident cases). Here, a parsimonious biomarker summary score was derived based on forward step-wise logistic models testing of all metabolites, using meta-analysis of the three derivation cohorts. Age, sex, baseline fasting glucose, and BMI were always included as covariates in the models. In each step, the metabolite with the lowest P-value was added as a covariate, and associations of all remaining metabolites with diabetes risk were assessed. This process was repeated until no further metabolites were significant at P<0.0009. The biomarker summary score was then defined as the sum of concentrations of the three selected metabolites weighted by beta-coefficients in the final stepwise model. The predictive ability of this biomarker summary score was then evaluated in NFBC, since this validation cohort had the highest number of cases and most reliably ascertained diagnoses. Odds ratios of the biomarker summary score were assessed both as a continuous marker and by quintiles, with adjustment for sex, baseline age, fasting glucose, and BMI. The influence of further adjusting for fasting insulin, triglycerides and HDL-cholesterol was also assessed. The potential improvement in risk discrimination when adding the biomarker summary score to models containing these two sets of clinical variables were compared in terms of C-statistic, integrated discrimination improvement and continuous reclassification (16). Statistical analyses were performed in R version 3.1.3.

## RESULTS

The study included 11,896 individuals from four Finnish cohorts. Characteristics of the study participants at the time of blood sampling are shown in **Table 1**. The mean age was 32.9 years (range 24-45). The follow-up time ranged from 8 to 15 years, during which a total of 392 incident cases of type 2 diabetes occurred. Mean concentrations and SDs of all metabolic measures are listed in **Supplementary Table 1**. The odds ratios of 104 selected metabolic measures with incident type 2 diabetes are shown in **Figure 1** and **Figure 2**; results for the remaining 125 metabolic measures assayed are found in **Supplementary Figure 2**. In meta-analysis of all four cohorts, 113 out of the 229 metabolic measures were robustly associated with incident type 2 diabetes (P<0.0009) when adjusting for sex, baseline age, BMI, and fasting glucose. The biomarkers associated with risk of future type 2 diabetes risk spanned multiple metabolic pathways of polar metabolites, fatty acids and detailed lipoprotein lipid measures, with significant odds ratios ranging from 1.18–1.50 for direct associations and from 0.59-0.86 for inverse associations per 1-SD metabolite concentration.

**Table 1.** Baseline characteristics of participants in the four prospective cohorts

|  | YFS | FINRISK-1997 | DILGOM | NFBC |
| --- | --- | --- | --- | --- |
| Number of individuals | 2141 | 3063 | 1421 | 5271 |
| Number of incident type 2 diabetes cases | 65 | 110 | 18 | 199 |
| Follow-up time (years) | 10 | 15 | 8 | 15 |
| Sex (% females) | 53.8 | 52.7 | 55.7 | 50.2 |
| Baseline Age (years) | 31.7 ± 4.7 | 35.3 ± 6.0 | 35.7 ± 6.2 | 31.2±0.4 |
| Body-mass index (kg/m<sup>2</sup>) | 25.0 ± 4.4 | 25.1 ± 4.2 | 25.6 ± 4.4 | 24.6 ± 4.1 |
| Glucose (mmol/l) | 5.0 ± 0.4 | 4.7 ± 0.6 | 5.6 ± 0.4 | 5.0 ± 0.4 |
| Total cholesterol (mmol/l) | 5.1 ± 1.0 | 5.1 ± 1.0 | 5.0 ± 0.9 | 5.0 ± 1.0 |
| HDL cholesterol (mmol/l) | 1.3 ± 0.3 | 1.4 ± 0.3 | 1.4 ± 0.4 | 1.5 ± 0.4 |
| Triglycerides (mmol/l) | 1.3 ± 0.8 | 1.3 ± 1.0 | 1.3 ± 0.9 | 1.2 ± 0.7 |
| Plasma Insulin (mU/l) | 7.6 ± 5.2 | 5.7 ± 5.5 | 5.6 ± 3.9 | 8.3 ± 3.9 |
| Lipid-lowering medication (%) | 0.3 | 0.3 | 1.3 | 0.1 |

Values are mean ± SD.
YFS, The Cardiovascular risk in Young Finns Study; DILGOM, Dietary, Lifestyle, and Genetic determinants of Obesity and Metabolic syndrome study; NFBC, Northern Finland Birth Cohort of 1966.

**Figure 1.**
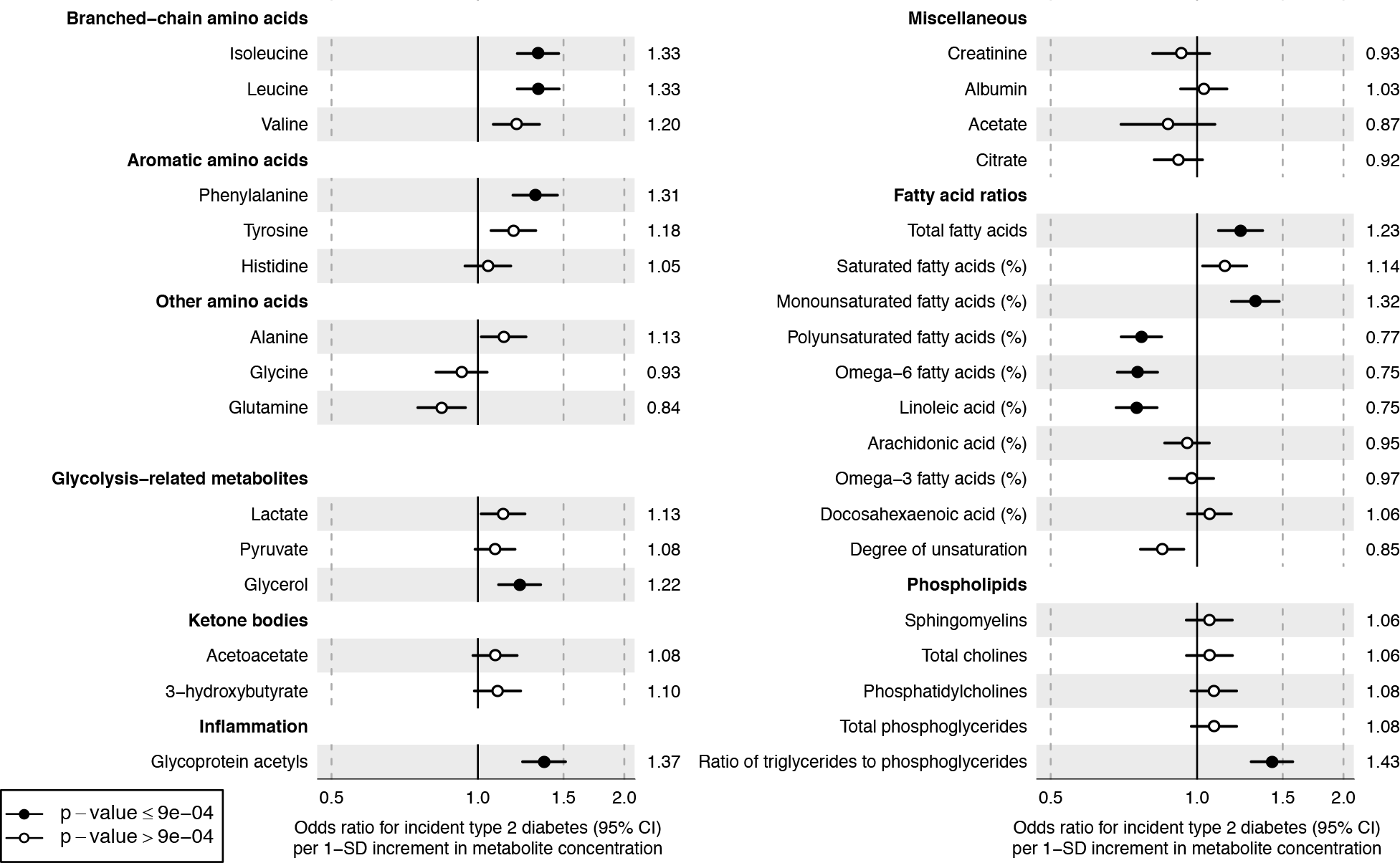
Relation of baseline circulating metabolite concentrations to risk of future type 2 diabetes. Values are odds ratios (95% confidence intervals) per 1-SD log-transformed metabolite concentration. Odds ratios were adjusted for sex, baseline age, body mass index and fasting glucose. The results were meta-analyzed for 11,896 young adults from four prospective cohorts.

**Figure 2.**
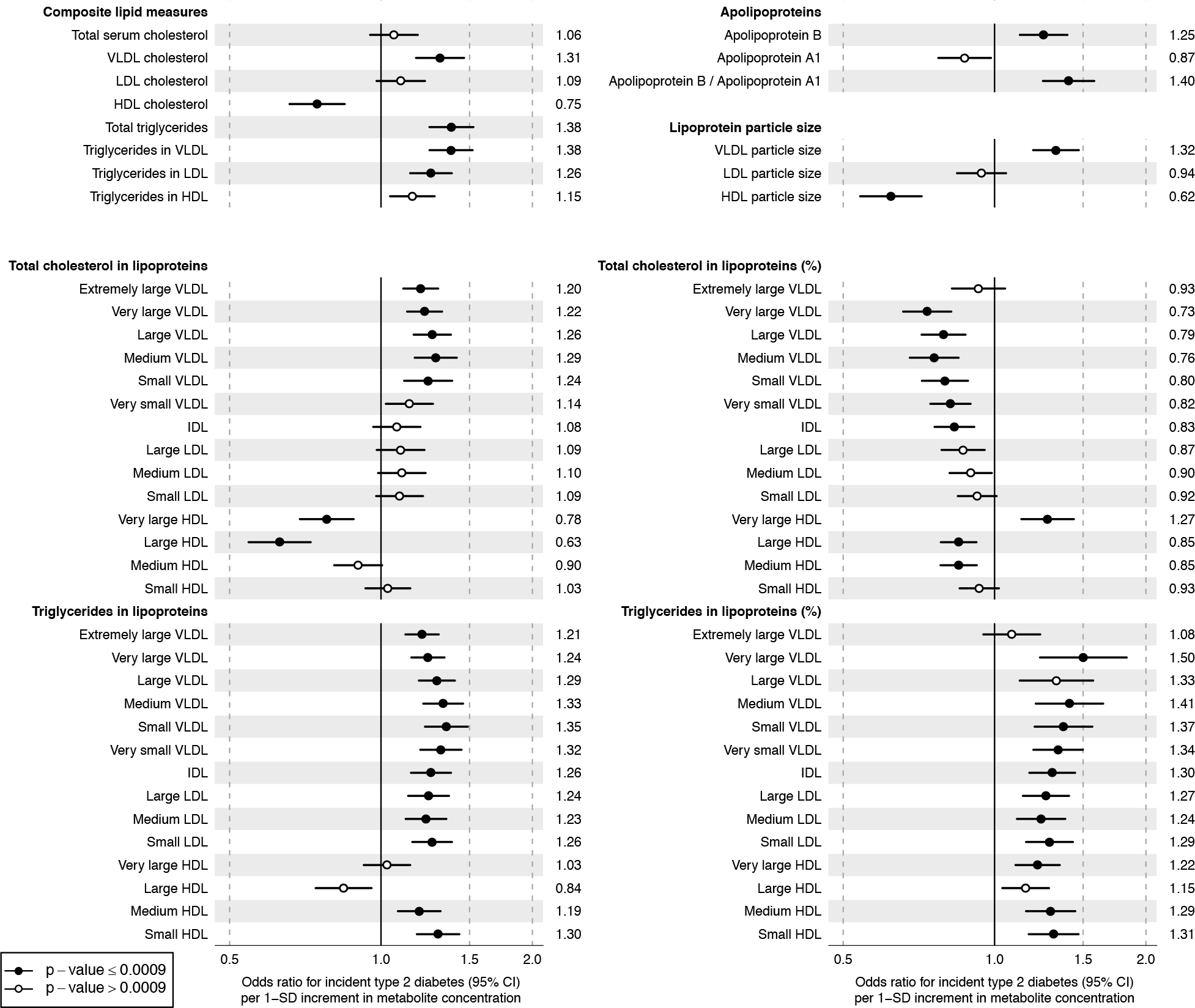
Relation of baseline circulating lipoprotein measures to risk of future type 2 diabetes. Values are odds ratios (95% confidence intervals) per 1-SD log-transformed metaboliteconcentration. Odds ratios were adjusted for sex, baseline age, body mass index and fasting glucose. The results were meta-analyzed for 11,896 young adults from four prospective cohorts. Odds ratios for the remaining 125 metabolic measures assayed are shown in Supplementary Figure 2.

### Amino acids, glycolysis, and inflammation

Branched-chain amino acids isoleucine, leucine, and valine (odds ratios 1.20–1.33) and the aromatic amino acids phenylalanine and tyrosine (1.33 and 1.18, respectively) were associated with the risk of type 2 diabetes (**Figure 1**). Glycerol was also associated with increased risk (1.22 [95% confidence interval 1.10–1.35]) while other glycolysis related metabolites had weaker associations. The inflammatory biomarker glycoprotein acetyls (GlycA) displayed one of the strongest associations for type 2 diabetes risk (1.37 [1.24–1.51]).

### Fatty acids

The total concentration of circulating fatty acids as well as the relative amount of monounsaturated fatty acids (MUFA; ratio to total fatty acids) were directly associated with increased risk for type 2 diabetes (1.32 [1.18–1.48]). In contrast, higher relative concentrations of omega-6 fatty acids were associated with decreased risk for type 2 diabetes (0.75 [0.69-0.83]). This inverse association was primarily driven by linoleic acid, whereas the association for arachidonic acid was weaker.

### Lipoprotein measures

Both lipid measures used in routine clinical settings and more fine-grained lipoprotein subclass measures are quantified by the NMR metabolomics platform. The associations of routine lipids, as well as cholesterol and triglyceride concentrations in 14 lipoprotein subclasses, with type 2 diabetes risk are shown in **Figure 2.** Additional lipoprotein subclass measures are shown in **Supplementary Figure 2**.

Overall, the cholesterol concentration within VLDL particles was associated with increased risk for type 2 diabetes, whereas the cholesterol in HDL particles was associated with decreased risk. Triglycerides in both VLDL, LDL, and HDL particles were strongly associated with increased diabetes risk. The analysis of 14 lipoprotein subclasses and their lipid content provides a more nuanced picture of these associations. Cholesterol in large HDL, in particular, was associated with decreased diabetes risk. The association patterns were similar for free cholesterol and cholesteryl esters; the strongest biomarker for decreased diabetes risk was free cholesterol in large HDL (0.59 [0.50-0.68]; **Supplementary Figure 2**). The absolute triglyceride concentrations within most of the lipoprotein subclasses were strongly associated with increased diabetes risk. This was also the case for the relative fraction of triglyceride within each subclass, i.e. the percentage of triglycerides to total lipids in a given lipoprotein particle size. Higher relative triglyceride content in lipoprotein particles generally reflects a lower cholesterol content, which is why inverse associations with future diabetes risk are seen for the percentage fractions of cholesterol in most of the lipoprotein subclasses.

Concentrations of apolipoproteins, the structural proteins of lipoprotein particles, were also associated with increased risk for type 2 diabetes. In particular, the ratio of apolipoprotein B to apolipoprotein A1 was among the strongest predictors (1.40 [1.25-1.58]). Further, larger VLDL particle size was associated with increased diabetes risk (1.32 [1.19-1.47]) whereas larger HDL particle size displayed an inverse association (0.62 [0.54-0.72]).

### Consistency across cohorts and influence of insulin adjustment

The association patterns between metabolites and incident type 2 diabetes were highly consistent in all four cohorts despite between-cohort differences in fasting status and ascertainment of diabetes diagnoses at follow-up (**Supplementary Figure 3**). The metabolite associations were also similar for men and women (**Supplementary Figure 4**).

Since the metabolite associations with type 2 diabetes risk in our study are reminiscent of those previously reported with glycemic traits and adiposity (17, 18), we examined the cross-sectional associations of metabolites with BMI, HOMA-IR index, and fasting glucose (**Supplementary Figure 5**). We found a high consistency in the overall pattern of metabolite associations for diabetes risk with those observed for BMI and HOMA-IR index, whereas the associations with fasting glucose were substantially weaker in these young adults.

To evaluate whether the metabolite biomarkers provide information beyond insulin resistance, we further adjusted analyses for baseline insulin levels. Most associations between metabolites and future risk of type 2 diabetes were moderately attenuated when including insulin as covariate, but the overall pattern persisted and 71 of the metabolic measures remained significant at P<0.0009 (**Supplementary Figure 6**).

### Biomarker score identifies individuals with increased risk of diabetes

To examine potential predictive utility of a combination of metabolic measures, we derived a biomarker summary score for the risk of future type 2 diabetes. Using a stepwise modeling approach based on three of the cohorts, three metabolic measures were selected as independent predictors of diabetes: phenylalanine, free cholesterol in large HDL, and cholesteryl esters (CE) to total lipids ratio within large VLDL. The biomarker summary score was then evaluated in the NFBC validation cohort. Here, the odds ratio of incident type 2 diabetes was 10.1 [4.2-24.1] among individuals in the upper fifth of the biomarker summary score compared to those in the lower fifth, when adjusting for age, sex, and baseline glucose and BMI (**Table 2**). If further adjusting for baseline insulin, triglycerides, and HDL-cholesterol, the odds ratio for individuals in the highest vs. lowest fifth of the biomarker summary score was attenuated to 5.79 [2.22-15.1]. The discrimination in absolute risk for future type 2 diabetes is presented in **Supplementary Tables 3** and **4** and **Supplementary Figure 7**; the prediction model enhanced with the biomarker summary score displayed improved risk discrimination above the basic clinical model (age, sex, BMI and fasting glucose), but only modest improvements above a model extended with triglycerides, HDL cholesterol, and fasting insulin.

**Table 2.** Biomarker summary score for the risk of type 2 diabetes during 15-year follow-up, assessed for 5,271 individuals aged 31 at blood sampling

| Individuals divided into fifths of biomarker summary score | Number of incident type 2 diabetes cases | Model 1: Age, sex, BMI and fasting glucose | Model 2: Model 1 + triglycerides, HDL-cholesterol and insulin as additional covariates |
| --- | --- | --- | --- |
|  | N (%) | Odds ratio (95% CI) | Odds ratio (95% CI) |
| Lower fifth | 6 (0.6) | Reference | Reference |
| 20% to 40% | 14 (1.3) | 2.17 (0.83-5.67) | 1.95 (0.74-5.16) |
| 40% to 60% | 36 (3.4) | 4.09 (2.08-12.0) | 3.93 (1.59-9.70) |
| 60% to 80% | 47 (4.5) | 5.92 (2.47-14.2) | 4.11 (1.63-10.3) |
| Upper fifth | 96 (9.1) | 10.1 (4.20-24.1) | 5.80 (2.22-15.1) |
| Per 1-SD increment |  | 1.76 (1.48-2.09)<br>P=2×10<sup>-10</sup> | 1.42 (1.14-1.76)<br>P=0.002 |

The biomarker summary score was calculated as the weighted sum of concentrations of three circulating metabolites: phenylalanine (weight = 0.320), free cholesterol in large HDL (weight = -0.474), and ratio of cholesteryl esters to total lipids in large VLDL (weight = -0.321). The beta-coefficients used as weights for the biomarkers score were derived by meta-analysis of three derivation cohorts.

## DISCUSSION

This large, multi-cohort study describes a metabolic signature of increased risk for type 2 diabetes in young adults up to 15 years prior to disease onset. Metabolic aberrations related to incident type 2 diabetes risk spanned amino acids, fatty acid balance, inflammation, and detailed lipoprotein particle composition, with consistent results across the four cohorts. Among the strongest biomarkers were higher concentrations of branched-chained and aromatic amino acids, VLDL particle measures, and the enrichment of triglycerides in all lipoprotein subclasses. Moreover, higher circulating levels of GlycA, glycerol, and MUFA also predicted increased risk for type 2 diabetes, whereas glutamine, linoleic acid, as well as HDL particle size and certain lipid measures within large HDL were associated with lower risk. A biomarker score consisting of three metabolic measures was predictive of a 10-fold elevation in the long-term risk for type 2 diabetes among 31-year old men and women.

The metabolic signature for type 2 diabetes risk described here included biomarkers across multiple molecular pathways. Branched-chain and aromatic amino acids were among the first biomarkers for type 2 diabetes risk identified by metabolomics (10). Their association with diabetes risk has since been replicated in several epidemiological studies (8, 9, 19) and extended to insulin resistance and glycemia (9, 18, 20). The odds ratios of all amino acids assayed in this study were consistent with a recent meta-analysis of prospective studies (8). We extend these prior findings by showing that branched-chain and aromatic amino acid levels are predictive of the long-term risk of type 2 diabetes already in young adults.

The causal relation between amino acid levels and type 2 diabetes risk is not yet fully clear. Mendelian randomization studies have indicated that adiposity and insulin resistance leads to increased branched-chain amino acid levels (12, 17), but also that the metabolism of these amino acids play a causal role in the development of type 2 diabetes (11). In addition, physiological studies have suggested mechanisms by which alterations in BCAA metabolism might cause insulin resistance and impairment of insulin secretion (21, 22). Altered amino acid metabolism may also represent a link between diabetes and cardiovascular diseases, common complications of type 2 diabetes (23, 24). Our results in young adults support the notion that amino acid profiling may prove helpful in monitoring of cardiometabolic health in asymptomatic individuals, with potential to facilitate targeted interventions and tracking of their effects (25).

Pervasive alterations in the lipoprotein profile were found to be predictive of future diabetes risk. These included both established lipid measures and novel associations of increased particle size and lipid content of VLDL, and enrichment of triglycerides in all lipoprotein subclasses. These lipid modulations are similar to those previously reported in cross-sectional settings for individuals with impaired glucose tolerance (26, 27). They may reflect early stages of the aberrations in lipoprotein metabolism characteristic of insulin resistance: increased production of large VLDLs, increased catabolism of HDLs, and increased transfer of triglycerides to HDL and LDL particles (28). Our findings indicate that such distortions of lipoprotein metabolism may already be present in normoglycemic young adults and reflect an increased risk for type 2 diabetes.

Increasing evidence suggests that certain fatty acid levels are predictive of type 2 diabetes. Our finding that a higher relative concentration of omega-6 fatty acids was associated with decreased diabetes risk, whereas higher MUFA levels were associated with increased diabetes risk is consistent with previous investigations (29, 30). Consistent with our results in young adults, a recent large-scale study pooling data from 20 prospective cohorts reported that higher levels of linoleic acid, the predominant omega-6 fatty acid, has long-term benefits for the prevention of type 2 diabetes (30). These fatty acids are likely reflective of both dietary composition and endogenous metabolism (31). Dietary counseling aiming to replace saturated fat with unsaturated fat in the diet, in accordance with Nordic dietary recommendations, has been shown to decrease circulating MUFA and increase circulating omega-3 and omega-6 levels (32). If these fatty acids play a causal role in the development of type 2 diabetes, then our results suggest that interventions modifying dietary fatty acid composition could be effective in prevention.

In addition to modulations in lipoprotein metabolism, metabolic measures related to lipolysis (glycerol) and inflammation (GlycA; a marker of chronic inflammation (33, 34)) were predictive biomarkers, illustrating how many different pathways are perturbed long before onset of type 2 diabetes. The overall metabolic signature of increased diabetes risk was reminiscent of the patterns of metabolite associations for adiposity and insulin resistance index. This is keeping with previous large-scale metabolic profiling studies (17, 18, 20) and consistent with the pathophysiology of type 2 diabetes, where insulin sensitivity gradually declines years before clinical disease onset (35). The similarity of the metabolic signatures for type 2 diabetes risk to those for higher BMI and insulin levels could suggests that the metabolic biomarkers for diabetes risk are manifestations of developing insulin resistance. Nonetheless, the overall pattern of biomarker associations remained predictive after controlling for baseline BMI and insulin levels. This indicates that comprehensive metabolic profiling is sensitive to detect subtle metabolic changes that precede insulin resistance and hyperglycemia in apparently healthy young adults.

Whereas the comprehensive signature of single-biomarkers for type 2 diabetes risk provide a picture of the numerous metabolic pathways reflective of the disease development, the measurement of multiple biomarkers in one go may prove beneficial for disease prediction. Supporting this notion, we found that a simple biomarker summary score comprised of phenylalanine and two detailed lipoprotein measures was a stronger predictor of diabetes risk than any of the individual biomarkers. The 10-fold elevation in diabetes risk observed for those in the highest compared to lowest fifth of the biomarker summary score indicates that multi-metabolite scores could potentially aid identification of high-risk individuals at young age. In this age group, the absolute risk for type 2 diabetes is low and a focus on relative risk compared to peers may be most effective for targeted prevention efforts. Future studies are needed to evaluate the potential of such scores for risk identification and health tracking in clinical settings.

Our study has both strengths and limitations. Strengths include the large sample size, and the profiling of multiple prospective cohorts. Our results were consistent across cohorts despite differences in age distribution, fasting status, and diagnostic ascertainment. The multi-cohort design allowed derivation of the biomarker summary score and validation in an independent cohort. Some limitations also need to be considered. First, because type 2 diabetes is relatively rare among young adults, the number of cases was modest despite the large sample size. The power for evaluating the predictive value of the biomarker scores was therefore limited. Second, since all cohorts were Finnish, our results cannot necessarily be generalized beyond white Europeans. However, previous research shows that amino acid measures may be even stronger predictors of type 2 diabetes in South Asians as compared to Europeans (36). Third, the NMR metabolomics platform is not able to quantify metabolites present in blood in very low concentrations, and we could therefore not replicate several previously reported metabolomic biomarkers for diabetes (8, 9, 37). Nonetheless, the NMR metabolomics method is high-throughput and consistent over time, and therefore particularly suited for large cohorts.

In conclusion, we have described a metabolic signature of increased risk for future type 2 diabetes in population-based cohorts of young adults with long follow-up. Metabolic aberrations were observed across multiple biological pathways, including inflammation, fatty acid balance, and aspects of lipoprotein metabolism. Our results extend the evidence of amino acid alterations as strong predictors of type 2 diabetes to young adults. If branched-chain amino acids, MUFAs or omega-6 fatty acids are proven to be causal in the pathogenesis of type 2 diabetes, then interventions aimed at altering the circulating levels may be beneficial already in early adulthood. Finally, we derived and validated a biomarker summary score that stratified the risk for future type 2 diabetes beyond risk factors used in primary care settings. These results support the possibility that early screening by detailed metabolic profiling could help targeting early interventions for type 2 diabetes prevention.

