## Supplementary Methods, Tables and Figures for "Circulating metabolites and the risk of type 2 diabetes: a prospective study of 11,896 young adults from four Finnish cohorts"

**Supplementary Methods: Cohort descriptions and type 2 diabetes status assessment.**
**Supplemental Table 1: Mean (SD) concentrations of 229 metabolic metabolites analyzed.**
**Supplemental Table 2: Tabulation of all biomarker results in the figures (online spreadsheet).**
**Supplemental Table 3: Regression models for risk of future type 2 diabetes.**
**Supplemental Table 4. Risk discrimination of three prediction models for risk of type 2 diabetes, assessed among 5,271 individuals aged 31.**
**Supplementary Figure 1: Overview of study cohorts and participant inclusion.**
**Supplementary Figure 2: Relation of 125 metabolic measures (not shown in main paper) to risk of future type 2 diabetes.**
**Supplementary Figure 3: Consistency of biomarkers to the risk of future diabetes across the four cohorts.**
**Supplementary Figure 4: Biomarkers for the risk of future diabetes assessed separately for men and women.**
**Supplementary Figure 5. Biomarkers for the risk of future diabetes compared to cross-sectional associations with BMI, HOMA-IR and fasting glucose.**
**Supplementary Figure 6: Biomarker associations with diabetes risk after additional adjustment for insulin and without adjustment for BMI.**
**Supplementary Figure 7. Refinement of absolute risk for type 2 diabetes by including biomarker summary score into prediction model.**

**Supplementary Methods: Cohort descriptions and type 2 diabetes status assessment.**

*The Cardiovascular Risk in Young Finns Study*

The Cardiovascular Risk in Young Finns Study (YFS) is an ongoing longitudinal multi-center follow-up of 3,596 Finnish individuals randomly chosen from the cities of Turku, Tampere, Helsinki, Oulu, and Kuopio, in addition to their rural surroundings (http://youngfinnsstudy.utu.fi)(1). The first cross-sectional survey was conducted in 1980 and the follow-up surveys concerning whole study population were conducted in 1983, 1986, 2001, 2007, and 2011. NMR metabolomics has been performed from n~2,200 serum samples drawn from all participants attending the 2001, 2007, and 2011 follow-up visits after overnight fasting using the Nightingale Health platform (2016 quantification version) (2). The metabolic measures acquired from serum samples from the 2001 collection (n=2,246 available) form the baseline data for the present study. Pregnant women were omitted from the analyses (n=78). Individuals with prevalent diabetes at the 2001-baseline were excluded from the analyses, based on self-reported diabetes (type 1 and type 2), use of anti-diabetic medications and fasting plasma glucose ≥ 7.0 mmol/l (n=23).

Outcome information on diabetes status at follow-up was based on a combination of fasting plasma glucose ≥7 mmol/L at either of the 2007 and the 2011 surveys, or if they reported having been given a type 2 diabetes diagnosis by a physician. Individuals whose $HbA_{1c}$ was ≥6.5% at the 2011 follow-up or who reported taking glucose-lowering medication at the 2007 or 2011 follow-up were also classified as having type 2 diabetes. To complement the information on diabetes outcome status from the follow-up surveys, type 2 diabetes diagnoses were obtained from the National Social Insurance Institution Drug Reimbursement Registry and nationwide hospital discharge registries, as detailed in the paragraph of registry tracking of diabetes incidence (3).

All participants gave written informed consent, and the study was approved by the local ethics committees of the study sites.

*FINRISK-1997*

FINRISK is a series of population-based health examination surveys to evaluate the cardiovascular risk status in Finnish population (4). The first survey was done in 1972 in provinces of North Karelia and Northern Savo. Subsequently, the FINRISK surveys have been expanded to surrounding regions of five Finnish cities: Turku, Loimaa, Helsinki, Vantaa, and Oulu. The surveys have been undertaken every five years and a new randomly chosen sample representative of population aged 25-74 (since 1997) has been invited to the surveys every time. Each study visit has contained questionnaires, physical examination and venous blood samples. Data for the present study stems from the FINRISK 1997 survey, in which 8,444 individuals participated. Serum samples for NMR metabolomics were available for 7,602 individuals, and measured using the Nightingale Health platform (2016 quantification version)(2). The median fasting time before the blood samples were drawn was 5 hours (interquartile range 4-6 hours). Individuals over 45 years at baseline were excluded from the analysis; out of 3516 participants aged 24–45, metabolomics data were available for 3210.

We used several data sources to ascertain exclusion of prevalent diabetes at baseline for (n=75): a) self-report of doctor-diagnosed diabetes or impaired glucose tolerance in the questionnaire, b) blood glucose ≥7.0 mmol/L at baseline, and c) the national drug reimbursement records and the National Hospital Discharge Register were checked for reimbursements of purchases of hypoglycemic drugs or hospitalizations with diabetes as the main or an additional diagnosis. If any of these sources was positive, the person was considered as having prevalent diabetes and was excluded from the analyses.

For tracking information about incidence of type 2 diabetes, we used information on National Social Insurance Institution Drug Reimbursement Registry and nation-wide hospital discharge register as described previously and detailed further below (5). The register data used in this study

included diagnoses made between spring 1997 and December 2012 (15-year follow-up). No laboratory data were available for FINRISK1997 to confirm incident diabetes status.
All participants gave written informed consent and the FINRISK study was approved by the ethical committee of the National Public Health Institute, Helsinki, Finland.

*Dietary, Lifestyle, and Genetic determinants of Obesity and Metabolic syndrome study*

The Dietary, Lifestyle, and Genetic determinants of Obesity and Metabolic syndrome (DILGOM) study is an extension of the FINRISK 2007 survey, focused on obesity and metabolic syndrome (6). All FINRISK 2007 participants were invited for a re-survey, and 84% participated(n=5,024). DILGOM study participants living in Southwest Finland were re-examined in 2014. They completed questionnaires, gave blood samples and underwent physical examination. Individuals living in the Oulu province, North Karelia, or Northern Savo completed a questionnaire but did not participate in physical examination nor gave blood samples.
In the present study, NMR metabolomics data from 4,816 individuals were available from fasting blood samples. Individuals over 45 years at baseline and pregnant women were excluded from the analysis, leaving 1,488 individuals for the analyses. Individuals with prevalent diabetes in the 2007 survey were excluded from the analyses, based on self-reported diabetes, use of anti-diabetic medications, and fasting plasma glucose ≥ 7.0 mmol/l or 2h glucose ≥11.0 (n=67). This information was further complemented from the nationwide drug-reimbursement and hospital registries as for FINRISK 1997 participants.
For incident diabetes, we used register-based diagnosis as described above for FINRISK 1997, follow-up time until end of 2014 (7.8 years). In addition, for participants living in southwestern Finland we had data fasting glucose available at the 7-year re-survey and individuals with fasting glucose ≥ 7.0 mmol/l were assigned as diabetes cases.
The Coordinating Ethics Committee of the Hospital District of Helsinki and Uusimaa has approved the FINRISK 2007 study and the DILGOM extension.

*Northern Finland Birth Cohort of 1966*

The Northern Finland Birth Cohort (NFBC) of 1966 is a longitudinal follow-up study of 12,058 children born into the cohort, comprising 96% of all births during 1966 in the region (www.oulu.fi/NFBC) (7). The participants were enrolled to the study in 1965 when their mothers were on their 24th gestational week. Majority of the children were born in 1966, excluding a small subset born in the end of 1965 or early 1967. Follow-up surveys have been arranged when the participants were 0-1, 14, 31 or 46 years of age. Data collection in 1997 included clinical examination and serum sampling at age 30–32 for 6,007 individuals, which form the baseline for the present study(2, 8). Attendees in the 1997 field study (70% of those invited for survey; 52% of original cohort) were representative of the original cohort.
NMR-based metabolomics (Nightingale Health Ltd, 2016 quantification version) was measured from 5,709 individuals with serum sample available, of which 96% were fasting samples. Individuals with prevalent diabetes at baseline were excluded from analyses. Information on diabetes outcome status was based on fasting glucose (≥ 7.0 mmol/l) and OGTT (2-hour value ≥ 11.0 mmol/l) assessed at the 2012 follow-up when the participants were 46 years old (data available for 3306 individuals). The diabetes outcome status from OGTT was complemented using nationwide reimbursement registries on prescription medication and hospital registry data for all participants with metabolomics at 31-year baseline as detailed below.
Informed written consent was obtained from all participants. The research protocols were approved by the Ethics Committees s of University of Oulu and that of Northern Ostrobotnia Hospital District, Finland.

*Registry information on incidence of type 2 diabetes*

Three data sources were used to identify cases of incident diabetes during the follow-up: 1) Record linkage of the study participants with the National Social Insurance Institution Drug Reimbursement Registry on the basis of the personal identification code unique to each individual in the country. The Social Insurance Institute keeps a nation-wide register of persons entitled to these reimbursements. 2) Record linkage with the National Hospital Discharge Register, which includes all hospitalizations in Finland (main diagnosis and up to four additional diagnoses). We checked whether diabetes (ICD-10 code E10-E14) was listed as any of the diagnoses for a hospitalization during the follow-up. 3) Record linkage with the National Causes-of-Death Register, which includes all deaths of permanent residents of Finland. We checked whether diabetes (ICD-10 code E10-E14) was mentioned as any of the causes of death (underlying cause of death, direct cause of death, or the contributing causes of death). If diabetes was found in any of these data sources, the person was considered to have incident diabetes. The date when the diabetes diagnosis first appeared was taken as the date of onset of diabetes. These procedures identify all cases of diabetes that were treated with hypoglycemic medications or hospitalized or who died during the follow-up. However, diabetic patients treated with diet only, who were not hospitalized and did not die, were not identified by these procedures (5).

**Supplementary Table 1. Mean (SD) concentrations of 229 metabolic measure analyzed and their odds ratios for incident type 2 diabetes**

| Metabolic measure | Units | Mean concen-tration | Standard deviation (SD) | Odds ratio per 1-SD | 95% CI | P-value |
|---|---|---|---|---|---|---|
| Isoleucine | mmol/l | 0.056 | 0.018 | 1.329 | 1.205–1.466 | 1E-8 |
| Leucine | mmol/l | 0.084 | 0.021 | 1.331 | 1.205–1.469 | 2E-8 |
| Valine | mmol/l | 0.20 | 0.048 | 1.200 | 1.076–1.339 | 0.001 |
| Phenylalanine | mmol/l | 0.081 | 0.013 | 1.312 | 1.180–1.458 | 5E-7 |
| Tyrosine | mmol/l | 0.054 | 0.013 | 1.183 | 1.064–1.315 | 0.002 |
| Alanine | mmol/l | 0.41 | 0.068 | 1.130 | 1.016–1.256 | 0.02 |
| Glutamine | mmol/l | 0.51 | 0.084 | 0.842 | 0.752–0.943 | 0.003 |
| Glycine | mmol/l | 0.30 | 0.063 | 0.926 | 0.820–1.045 | 0.21 |
| Histidine | mmol/l | 0.071 | 0.011 | 1.048 | 0.941–1.168 | 0.39 |
| Lactate | mmol/l | 1.38 | 0.38 | 1.126 | 1.015–1.250 | 0.02 |
| Pyruvate | mmol/l | 0.083 | 0.027 | 1.084 | 0.987–1.192 | 0.09 |
| Glycerol | mmol/l | 0.11 | 0.058 | 1.218 | 1.103–1.346 | 1E-4 |
| Acetoacetate | mmol/l | 0.062 | 0.049 | 1.085 | 0.977–1.204 | 0.13 |
| beta-hydroxybutyrate | mmol/l | 0.18 | 0.14 | 1.097 | 0.983–1.224 | 0.10 |
| Glycoprotein Acetyls | mmol/l | 1.34 | 0.23 | 1.368 | 1.236–1.514 | 2E-9 |
| Creatinine | mmol/l | 0.062 | 0.013 | 0.927 | 0.810–1.061 | 0.27 |
| Albumin | signal area | 0.10 | 0.011 | 1.032 | 0.925–1.152 | 0.57 |
| Acetate | mmol/l | 0.048 | 0.033 | 0.871 | 0.697–1.088 | 0.22 |
| Citrate | mmol/l | 0.11 | 0.019 | 0.915 | 0.816–1.026 | 0.13 |
| Total fatty acids | mmol/l | 12.03 | 2.95 | 1.229 | 1.107–1.364 | 1E-4 |
| Saturated fatty acids, relative to total fatty acids | % | 36.90 | 2.29 | 1.140 | 1.027–1.265 | 0.01 |
| Monounsaturated fatty acids, relative to total fatty acids | % | 25.60 | 2.86 | 1.317 | 1.176–1.475 | 2E-6 |
| Polyunsaturated fatty acids, relative to total fatty acids | % | 37.52 | 3.62 | 0.768 | 0.698–0.845 | 7E-8 |
| Omega-6 fatty acids, relative to total fatty acids | % | 33.02 | 3.24 | 0.754 | 0.686–0.829 | 6E-9 |
| Linoleic acid, relative to total fatty acids | % | 26.95 | 3.52 | 0.751 | 0.681–0.828 | 7E-9 |
| Arachidonic acid, relative to total fatty acids | % | 6.07 | 1.13 | 0.953 | 0.858–1.060 | 0.38 |
| Omega-3 fatty acids, relative to total fatty acids | % | 4.49 | 1.09 | 0.974 | 0.878–1.082 | 0.63 |
| Docosahexaenoic acid, relative to total fatty acids | % | 1.60 | 0.56 | 1.060 | 0.956–1.176 | 0.27 |
| Unsaturation degree | - | 1.19 | 0.061 | 0.847 | 0.765–0.939 | 0.002 |
| Sphingomyelins | mmol/l | 0.52 | 0.10 | 1.060 | 0.950–1.181 | 0.30 |
| Total cholines | mmol/l | 2.49 | 0.51 | 1.060 | 0.951–1.182 | 0.29 |
| Phosphatidylcholines | mmol/l | 2.05 | 0.44 | 1.082 | 0.971–1.206 | 0.15 |
| Total phosphoglycerides | mmol/l | 2.05 | 0.48 | 1.083 | 0.973–1.206 | 0.14 |

| | | | | | | |
|---|---|---|---|---|---|---|
| Ratio of triglycerides to phosphoglycerides | - | 0.56 | 0.21 | 1.425 | 1.292–1.573 | 2E-12 |
| Total serum cholesterol | mmol/l | 4.82 | 1.07 | 1.061 | 0.951–1.183 | 0.29 |
| VLDL cholesterol | mol/l | 0.76 | 0.33 | 1.310 | 1.174–1.463 | 1E-6 |
| LDL cholesterol | mmol/l | 1.73 | 0.57 | 1.095 | 0.980–1.223 | 0.11 |
| HDL cholesterol | mmol/l | 1.56 | 0.38 | 0.747 | 0.658–0.847 | 6E-6 |
| Total triglycerides | mmol/l | 1.16 | 0.62 | 1.381 | 1.249–1.527 | 3E-10 |
| VLDL triglycerides | mmol/l | 0.70 | 0.49 | 1.378 | 1.249–1.521 | 2E-10 |
| LDL triglycerides | mmol/l | 0.19 | 0.10 | 1.257 | 1.143–1.383 | 3E-6 |
| HDL triglycerides | mmol/l | 0.15 | 0.05 | 1.154 | 1.042–1.279 | 0.006 |
| Apolipoprotein B | g/l | 0.90 | 0.23 | 1.252 | 1.123–1.396 | 5E-5 |
| Apolipoprotein A1 | g/l | 1.59 | 0.23 | 0.872 | 0.773–0.984 | 0.03 |
| ApoB ratio to ApoA1 | - | 0.58 | 0.16 | 1.403 | 1.248–1.578 | 2E-8 |
| VLDL particle size, diameter | nm | 35.79 | 1.33 | 1.325 | 1.193–1.471 | 1E-7 |
| LDL particle size, diameter | nm | 23.66 | 0.17 | 0.942 | 0.842–1.054 | 0.30 |
| HDL particle size, diameter | nm | 10.07 | 0.30 | 0.622 | 0.541–0.716 | 4E-11 |
| XXL VLDL cholesterol | mmol/l | 0.005 | 0.006 | 1.200 | 1.108–1.299 | 8E-6 |
| XL VLDL cholesterol | mmol/l | 0.013 | 0.016 | 1.221 | 1.127–1.323 | 1E-6 |
| L VLDL cholesterol | mmol/l | 0.051 | 0.053 | 1.265 | 1.161–1.377 | 7E-8 |
| M VLDL cholesterol | mmol/l | 0.15 | 0.09 | 1.285 | 1.167–1.415 | 3E-7 |
| S VLDL cholesterol | mmol/l | 0.24 | 0.10 | 1.241 | 1.112–1.385 | 1E-4 |
| XS VLDL cholesterol | mmol/l | 0.30 | 0.10 | 1.139 | 1.023–1.269 | 0.02 |
| IDL cholesterol | mmol/l | 0.77 | 0.23 | 1.075 | 0.965–1.198 | 0.19 |
| L LDL cholesterol | mmol/l | 0.92 | 0.29 | 1.094 | 0.981–1.220 | 0.11 |
| M LDL cholesterol | mmol/l | 0.50 | 0.18 | 1.100 | 0.987–1.226 | 0.09 |
| S LDL cholesterol | mmol/l | 0.31 | 0.11 | 1.089 | 0.978–1.212 | 0.12 |
| XL HDL cholesterol | mmol/l | 0.28 | 0.15 | 0.780 | 0.690–0.882 | 8E-5 |
| L HDL cholesterol | mmol/l | 0.41 | 0.21 | 0.629 | 0.545–0.725 | 2E-10 |
| M HDL cholesterol | mmol/l | 0.43 | 0.15 | 0.901 | 0.807–1.005 | 0.06 |
| S HDL cholesterol | mmol/l | 0.45 | 0.10 | 1.032 | 0.931–1.144 | 0.55 |
| XXL VLDL triglycerides | mmol/l | 0.018 | 0.020 | 1.207 | 1.118–1.302 | 1E-6 |
| XL VLDL triglycerides | mmol/l | 0.030 | 0.041 | 1.240 | 1.149–1.337 | 3E-8 |
| L VLDL triglycerides | mmol/l | 0.10 | 0.12 | 1.292 | 1.189–1.403 | 1E-9 |
| M VLDL triglycerides | mmol/l | 0.22 | 0.17 | 1.329 | 1.214–1.456 | 9E-10 |
| S VLDL triglycerides | mmol/l | 0.21 | 0.11 | 1.349 | 1.223–1.488 | 2E-9 |
| XS VLDL triglycerides | mmol/l | 0.11 | 0.05 | 1.316 | 1.197–1.446 | 1E-8 |
| IDL triglycerides | mmol/l | 0.12 | 0.06 | 1.257 | 1.147–1.378 | 1E-6 |
| L LDL triglycerides | mmol/l | 0.11 | 0.05 | 1.243 | 1.132–1.365 | 5E-6 |
| M LDL triglycerides | mmol/l | 0.051 | 0.028 | 1.229 | 1.120–1.349 | 1E-5 |
| S LDL triglycerides | mmol/l | 0.031 | 0.017 | 1.264 | 1.154–1.384 | 4E-7 |
| XL HDL triglycerides | mmol/l | 0.017 | 0.010 | 1.028 | 0.924–1.143 | 0.61 |
| L HDL triglycerides | mmol/l | 0.036 | 0.019 | 0.843 | 0.741–0.957 | 0.009 |

| | | | | | | |
|---|---|---|---|---|---|---|
| M HDL triglycerides | mmol/l | 0.045 | 0.018 | 1.191 | 1.081–1.313 | 4E-4 |
| S HDL triglycerides | mmol/l | 0.049 | 0.020 | 1.299 | 1.179–1.432 | 1E-7 |
| XXL VLDL cholesterol, ratio to total lipids | % | 19.04 | 5.21 | 0.929 | 0.823–1.049 | 0.24 |
| XL VLDL cholesterol, ratio to total lipids | % | 29.13 | 11.80 | 0.735 | 0.658–0.820 | 4E-8 |
| L VLDL cholesterol, ratio to total lipids | % | 28.23 | 8.26 | 0.792 | 0.716–0.875 | 5E-6 |
| M VLDL cholesterol, ratio to total lipids | % | 32.97 | 6.47 | 0.759 | 0.678–0.848 | 1E-6 |
| S VLDL cholesterol, ratio to total lipids | % | 41.54 | 6.49 | 0.797 | 0.717–0.886 | 2E-5 |
| XS VLDL cholesterol, ratio to total lipids | % | 52.01 | 5.32 | 0.817 | 0.745–0.896 | 2E-5 |
| IDL cholesterol, ratio to total lipids | % | 63.14 | 3.10 | 0.832 | 0.760–0.911 | 7E-5 |
| L LDL cholesterol, ratio to total lipids | % | 66.63 | 3.06 | 0.866 | 0.784–0.956 | 0.004 |
| M LDL cholesterol, ratio to total lipids | % | 65.13 | 4.77 | 0.897 | 0.814–0.988 | 0.03 |
| S LDL cholesterol, ratio to total lipids | % | 62.16 | 5.08 | 0.924 | 0.845–1.009 | 0.08 |
| XL HDL cholesterol, ratio to total lipids | % | 48.95 | 8.30 | 1.275 | 1.130–1.438 | 8E-5 |
| L HDL cholesterol, ratio to total lipids | % | 47.20 | 5.08 | 0.848 | 0.782–0.921 | 8E-5 |
| M HDL cholesterol, ratio to total lipids | % | 47.30 | 5.30 | 0.848 | 0.781–0.922 | 1E-4 |
| S HDL cholesterol, ratio to total lipids | % | 41.00 | 4.49 | 0.932 | 0.852–1.020 | 0.13 |
| XXL VLDL triglycerides, ratio to total lipids | % | 70.67 | 6.17 | 1.082 | 0.950–1.233 | 0.24 |
| XL VLDL triglycerides, ratio to total lipids | % | 53.20 | 13.70 | 1.502 | 1.231–1.831 | 6E-5 |
| L VLDL triglycerides, ratio to total lipids | % | 52.98 | 8.98 | 1.327 | 1.122–1.570 | 0.001 |
| M VLDL triglycerides, ratio to total lipids | % | 45.73 | 7.17 | 1.409 | 1.208–1.644 | 1E-5 |
| S VLDL triglycerides, ratio to total lipids | % | 35.09 | 6.33 | 1.371 | 1.201–1.565 | 3E-6 |
| XS VLDL triglycerides, ratio to total lipids | % | 19.12 | 4.81 | 1.338 | 1.195–1.499 | 5E-7 |
| IDL triglycerides, ratio to total lipids | % | 10.13 | 3.07 | 1.302 | 1.173–1.446 | 8E-7 |
| L LDL triglycerides, ratio to total lipids | % | 8.03 | 2.77 | 1.265 | 1.139–1.406 | 1E-5 |
| M LDL triglycerides, ratio to total lipids | % | 6.72 | 2.81 | 1.237 | 1.108–1.382 | 2E-4 |
| S LDL triglycerides, ratio to total lipids | % | 6.27 | 2.64 | 1.286 | 1.155–1.432 | 4E-6 |
| XL HDL triglycerides, ratio to total lipids | % | 3.78 | 3.54 | 1.218 | 1.101–1.348 | 1E-4 |

| | | | | | | |
|---|---|---|---|---|---|---|
| L HDL triglycerides, ratio to total lipids | % | 4.51 | 2.37 | 1.153 | 1.036–1.283 | 0.009 |
| M HDL triglycerides, ratio to total lipids | % | 5.36 | 2.64 | 1.293 | 1.155–1.447 | 8E-6 |
| S HDL triglycerides, ratio to total lipids | % | 4.58 | 2.03 | 1.311 | 1.169–1.469 | 3E-6 |
| HDL2 cholesterol | mmol/l | 1.04 | 0.36 | 0.733 | 0.647–0.831 | 1E-6 |
| HDL3 cholesterol | mmol/l | 0.52 | 0.05 | 0.981 | 0.883–1.090 | 0.72 |
| Esterified cholesterol | mmol/l | 3.43 | 0.77 | 1.057 | 0.948–1.179 | 0.32 |
| Free cholesterol | mmol/l | 1.39 | 0.30 | 1.074 | 0.964–1.196 | 0.19 |
| Remnant cholesterol | mmol/l | 1.53 | 0.51 | 1.232 | 1.099–1.380 | 3E-4 |
| Saturated fatty acids | mmol/l | 4.46 | 1.22 | 1.241 | 1.121–1.375 | 3E-5 |
| Monounsaturated fatty acids | mmol/l | 3.12 | 1.06 | 1.290 | 1.164–1.429 | 1E-6 |
| Polyunsaturated fatty acids | mmol/l | 4.45 | 0.86 | 1.100 | 0.988–1.224 | 0.08 |
| Omega-6 fatty acids | mmol/l | 3.91 | 0.75 | 1.084 | 0.974–1.207 | 0.14 |
| Linoleic acid | mmol/l | 3.18 | 0.59 | 1.051 | 0.944–1.171 | 0.36 |
| Arachidonic acid | mmol/l | 0.73 | 0.23 | 1.190 | 1.072–1.320 | 0.001 |
| Omega-3 fatty acids | mmol/l | 0.54 | 0.17 | 1.136 | 1.029–1.254 | 0.01 |
| Docosahexaenoic acid | mmol/l | 0.19 | 0.07 | 1.174 | 1.066–1.292 | 0.001 |
| XXL VLDL lipid concentration | mmol/l | 0.027 | 0.029 | 1.211 | 1.121–1.309 | 1E-6 |
| XL VLDL lipid concentration | mmol/l | 0.052 | 0.067 | 1.250 | 1.155–1.353 | 3E-8 |
| L VLDL lipid concentration | mmol/l | 0.19 | 0.21 | 1.312 | 1.201–1.433 | 2E-9 |
| M VLDL lipid concentration | mmol/l | 0.47 | 0.32 | 1.348 | 1.221–1.488 | 3E-9 |
| S VLDL lipid concentration | mmol/l | 0.58 | 0.25 | 1.353 | 1.212–1.510 | 7E-8 |
| XS VLDL lipid concentration | mmol/l | 0.57 | 0.18 | 1.223 | 1.099–1.360 | 2E-4 |
| IDL lipid concentration | mmol/l | 1.21 | 0.34 | 1.119 | 1.004–1.247 | 0.04 |
| L LDL lipid concentration | mmol/l | 1.37 | 0.40 | 1.127 | 1.010–1.258 | 0.03 |
| M LDL lipid concentration | mmol/l | 0.76 | 0.24 | 1.143 | 1.025–1.275 | 0.02 |
| S LDL lipid concentration | mmol/l | 0.49 | 0.15 | 1.140 | 1.023–1.269 | 0.02 |
| XL HDL lipid concentration | mmol/l | 0.56 | 0.29 | 0.717 | 0.630–0.815 | 4E-7 |
| L HDL lipid concentration | mmol/l | 0.84 | 0.39 | 0.671 | 0.591–0.763 | 1E-9 |
| M HDL lipid concentration | mmol/l | 0.89 | 0.26 | 0.933 | 0.840–1.037 | 0.20 |
| S HDL lipid concentration | mmol/l | 1.09 | 0.18 | 1.076 | 0.973–1.189 | 0.15 |
| XXL VLDL particle concentration | mol/l | 1.24E-10 | 1.33E-10 | 1.194 | 1.106–1.288 | 5E-6 |
| XL VLDL particle concentration | mol/l | 5.20E-10 | 6.79E-10 | 1.217 | 1.129–1.312 | 3E-7 |
| L VLDL particle concentration | mol/l | 3.25E-9 | 3.63E-9 | 1.245 | 1.153–1.345 | 3E-8 |
| M VLDL particle concentration | mol/l | 1.39E-8 | 9.52E-9 | 1.277 | 1.174–1.390 | 1E-8 |
| S VLDL particle concentration | mol/l | 2.91E-8 | 1.29E-8 | 1.311 | 1.189–1.445 | 6E-8 |

| | | | | | | |
|---|---|---|---|---|---|---|
| XS VLDL particle concentration | mol/l | 4.44E-8 | 1.45E-8 | 1.218 | 1.104–1.344 | 9E-5 |
| IDL particle concentration | mol/l | 1.20E-7 | 3.37E-8 | 1.125 | 1.016–1.247 | 0.02 |
| L LDL particle concentration | mol/l | 1.92E-7 | 5.64E-8 | 1.132 | 1.021–1.254 | 0.02 |
| M LDL particle concentration | mol/l | 1.49E-7 | 4.81E-8 | 1.145 | 1.033–1.269 | 0.01 |
| S LDL particle concentration | mol/l | 1.72E-7 | 5.25E-8 | 1.146 | 1.034–1.270 | 0.009 |
| XL HDL particle concentration | mol/l | 5.47E-7 | 2.83E-7 | 0.677 | 0.585–0.783 | 2E-7 |
| L HDL particle concentration | mol/l | 1.34E-6 | 6.06E-7 | 0.646 | 0.556–0.751 | 1E-8 |
| M HDL particle concentration | mol/l | 2.09E-6 | 6.02E-7 | 0.959 | 0.860–1.069 | 0.45 |
| S HDL particle concentration | mol/l | 4.91E-6 | 8.12E-7 | 1.109 | 1.004–1.226 | 0.04 |
| XXL VLDL cholesteryl esters | mmol/l | 3.29E-3 | 3.43E-3 | 1.191 | 1.096–1.294 | 4E-5 |
| XL VLDL cholesteryl esters | mmol/l | 0.0072 | 0.0085 | 1.214 | 1.120–1.316 | 2E-6 |
| L VLDL cholesteryl esters | mmol/l | 0.030 | 0.028 | 1.247 | 1.142–1.361 | 9E-7 |
| M VLDL cholesteryl esters | mmol/l | 0.095 | 0.052 | 1.244 | 1.128–1.372 | 1E-5 |
| S VLDL cholesteryl esters | mmol/l | 0.16 | 0.069 | 1.185 | 1.061–1.322 | 0.002 |
| XS VLDL cholesteryl esters | mmol/l | 0.21 | 0.072 | 1.131 | 1.015–1.260 | 0.03 |
| IDL cholesteryl esters | mmol/l | 0.55 | 0.16 | 1.084 | 0.973–1.208 | 0.14 |
| L LDL cholesteryl esters | mmol/l | 0.65 | 0.22 | 1.105 | 0.991–1.233 | 0.07 |
| M LDL cholesteryl esters | mmol/l | 0.35 | 0.15 | 1.094 | 0.981–1.219 | 0.11 |
| S LDL cholesteryl esters | mmol/l | 0.21 | 0.09 | 1.080 | 0.971–1.202 | 0.16 |
| XL HDL cholesteryl esters | mmol/l | 0.20 | 0.11 | 0.787 | 0.697–0.889 | 1E-4 |
| L HDL cholesteryl esters | mmol/l | 0.32 | 0.16 | 0.636 | 0.551–0.733 | 5E-10 |
| M HDL cholesteryl esters | mmol/l | 0.34 | 0.12 | 0.900 | 0.807–1.005 | 0.06 |
| S HDL cholesteryl esters | mmol/l | 0.33 | 0.08 | 1.018 | 0.919–1.129 | 0.73 |
| XXL VLDL free cholesterol | mmol/l | 0.0020 | 0.0024 | 1.199 | 1.111–1.293 | 3E-6 |
| XL VLDL free cholesterol | mmol/l | 0.006 | 0.007 | 1.219 | 1.127–1.319 | 8E-7 |
| L VLDL free cholesterol | mmol/l | 0.021 | 0.026 | 1.256 | 1.160–1.359 | 2E-8 |
| M VLDL free cholesterol | mmol/l | 0.055 | 0.041 | 1.294 | 1.184–1.413 | 1E-8 |
| S VLDL free cholesterol | mmol/l | 0.082 | 0.036 | 1.297 | 1.170–1.436 | 7E-7 |
| XS VLDL free cholesterol | mmol/l | 0.091 | 0.030 | 1.140 | 1.029–1.263 | 0.01 |
| IDL free cholesterol | mmol/l | 0.22 | 0.07 | 1.050 | 0.945–1.166 | 0.37 |
| L LDL free cholesterol | mmol/l | 0.26 | 0.07 | 1.060 | 0.953–1.178 | 0.28 |
| M LDL free cholesterol | mmol/l | 0.15 | 0.04 | 1.121 | 1.009–1.246 | 0.03 |
| S LDL free cholesterol | mmol/l | 0.092 | 0.022 | 1.116 | 1.005–1.241 | 0.04 |
| XL HDL free cholesterol | mmol/l | 0.075 | 0.039 | 0.754 | 0.659–0.863 | 4E-5 |
| L HDL free cholesterol | mmol/l | 0.087 | 0.049 | 0.587 | 0.501–0.687 | 4E-11 |
| M HDL free cholesterol | mmol/l | 0.082 | 0.030 | 0.924 | 0.826–1.034 | 0.17 |
| S HDL free cholesterol | mmol/l | 0.116 | 0.022 | 1.072 | 0.970–1.186 | 0.17 |

| | | | | | | |
|---|---|---|---|---|---|---|
| XXL VLDL phospholipids | mmol/l | 0.0029 | 0.0035 | 1.184 | 1.096–1.279 | 2E-5 |
| XL VLDL phospholipids | mmol/l | 0.009 | 0.012 | 1.224 | 1.134–1.321 | 2E-7 |
| L VLDL phospholipids | mmol/l | 0.035 | 0.039 | 1.270 | 1.172–1.377 | 6E-9 |
| M VLDL phospholipids | mmol/l | 0.10 | 0.06 | 1.307 | 1.194–1.431 | 7E-9 |
| S VLDL phospholipids | mmol/l | 0.13 | 0.05 | 1.321 | 1.191–1.464 | 1E-7 |
| XS VLDL phospholipids | mmol/l | 0.16 | 0.05 | 1.158 | 1.046–1.281 | 0.005 |
| IDL phospholipids | mmol/l | 0.32 | 0.08 | 1.098 | 0.988–1.220 | 0.08 |
| L LDL phospholipids | mmol/l | 0.34 | 0.08 | 1.120 | 1.008–1.246 | 0.04 |
| M LDL phospholipids | mmol/l | 0.21 | 0.05 | 1.190 | 1.072–1.321 | 0.001 |
| S LDL phospholipids | mmol/l | 0.15 | 0.03 | 1.181 | 1.066–1.309 | 0.002 |
| XL HDL phospholipids | mmol/l | 0.26 | 0.14 | 0.635 | 0.549–0.733 | 7E-10 |
| L HDL phospholipids | mmol/l | 0.40 | 0.17 | 0.690 | 0.605–0.786 | 2E-8 |
| M HDL phospholipids | mmol/l | 0.41 | 0.11 | 0.957 | 0.860–1.064 | 0.41 |
| S HDL phospholipids | mmol/l | 0.59 | 0.11 | 1.065 | 0.964–1.176 | 0.21 |
| XXL VLDL cholesteryl esters, ratio to total lipids | % | 12.25 | 4.63 | 0.882 | 0.788–0.987 | 0.03 |
| XL VLDL cholesteryl esters, ratio to total lipids | % | 15.46 | 7.10 | 0.765 | 0.686–0.852 | 1E-6 |
| L VLDL cholesteryl esters, ratio to total lipids | % | 18.22 | 7.62 | 0.770 | 0.696–0.853 | 5E-7 |
| M VLDL cholesteryl esters, ratio to total lipids | % | 21.85 | 6.34 | 0.757 | 0.685–0.838 | 6E-8 |
| S VLDL cholesteryl esters, ratio to total lipids | % | 27.46 | 6.23 | 0.797 | 0.719–0.882 | 1E-5 |
| XS VLDL cholesteryl esters, ratio to total lipids | % | 36.09 | 5.33 | 0.851 | 0.775–0.934 | 7E-4 |
| IDL cholesteryl esters, ratio to total lipids | % | 45.15 | 2.76 | 0.901 | 0.820–0.990 | 0.03 |
| L LDL cholesteryl esters, ratio to total lipids | % | 47.16 | 3.26 | 0.962 | 0.856–1.080 | 0.51 |
| M LDL cholesteryl esters, ratio to total lipids | % | 44.88 | 6.55 | 0.969 | 0.876–1.071 | 0.54 |
| S LDL cholesteryl esters, ratio to total lipids | % | 42.89 | 6.58 | 0.970 | 0.885–1.064 | 0.52 |
| XL HDL cholesteryl esters, ratio to total lipids | % | 35.51 | 7.82 | 1.200 | 1.065–1.353 | 0.003 |
| L HDL cholesteryl esters, ratio to total lipids | % | 37.43 | 3.60 | 0.910 | 0.838–0.988 | 0.02 |
| M HDL cholesteryl esters, ratio to total lipids | % | 38.22 | 4.88 | 0.864 | 0.795–0.940 | 7E-4 |
| S HDL cholesteryl esters, ratio to total lipids | % | 30.35 | 5.06 | 0.931 | 0.854–1.014 | 0.10 |
| XXL VLDL free cholesterol, ratio to total lipids | % | 6.79 | 2.18 | 1.138 | 0.992–1.304 | 0.06 |
| XL VLDL free cholesterol, ratio to total lipids | % | 13.68 | 6.35 | 0.777 | 0.689–0.875 | 3E-5 |
| L VLDL free cholesterol, ratio to total lipids | % | 10.01 | 3.03 | 1.163 | 0.995–1.359 | 0.06 |
| M VLDL free cholesterol, ratio to total lipids | % | 11.12 | 1.65 | 1.219 | 1.044–1.422 | 0.01 |

| | | | | | | |
|---|---|---|---|---|---|---|
| S VLDL free cholesterol, ratio to total lipids | % | 14.09 | 0.87 | 0.915 | 0.827–1.011 | 0.08 |
| XS VLDL free cholesterol, ratio to total lipids | % | 15.92 | 1.11 | 0.864 | 0.799–0.936 | 3E-4 |
| IDL free cholesterol, ratio to total lipids | % | 17.99 | 1.14 | 0.887 | 0.825–0.954 | 0.001 |
| L LDL free cholesterol, ratio to total lipids | % | 19.48 | 1.11 | 0.812 | 0.748–0.882 | 7E-7 |
| M LDL free cholesterol, ratio to total lipids | % | 20.25 | 2.34 | 0.849 | 0.753–0.958 | 0.008 |
| S LDL free cholesterol, ratio to total lipids | % | 19.27 | 2.03 | 0.861 | 0.768–0.966 | 0.01 |
| XL HDL free cholesterol, ratio to total lipids | % | 13.45 | 2.14 | 1.216 | 1.084–1.364 | 9E-4 |
| L HDL free cholesterol, ratio to total lipids | % | 9.78 | 1.90 | 0.771 | 0.710–0.837 | 7E-10 |
| M HDL free cholesterol, ratio to total lipids | % | 9.08 | 0.82 | 0.860 | 0.794–0.930 | 2E-4 |
| S HDL free cholesterol, ratio to total lipids | % | 10.64 | 0.84 | 0.975 | 0.880–1.079 | 0.62 |
| XXL VLDL phospholipids, ratio to total lipids | % | 10.30 | 2.46 | 0.999 | 0.883–1.130 | 0.99 |
| XL VLDL phospholipids, ratio to total lipids | % | 17.67 | 4.33 | 0.892 | 0.791–1.006 | 0.06 |
| L VLDL phospholipids, ratio to total lipids | % | 18.80 | 2.50 | 0.986 | 0.874–1.113 | 0.82 |
| M VLDL phospholipids, ratio to total lipids | % | 21.32 | 1.30 | 0.778 | 0.683–0.885 | 1E-4 |
| S VLDL phospholipids, ratio to total lipids | % | 23.38 | 1.67 | 0.837 | 0.748–0.936 | 0.002 |
| XS VLDL phospholipids, ratio to total lipids | % | 28.89 | 2.93 | 0.917 | 0.828–1.016 | 0.10 |
| IDL phospholipids, ratio to total lipids | % | 26.74 | 1.09 | 0.842 | 0.758–0.934 | 0.001 |
| L LDL phospholipids, ratio to total lipids | % | 25.34 | 1.69 | 0.905 | 0.807–1.014 | 0.09 |
| M LDL phospholipids, ratio to total lipids | % | 28.16 | 3.41 | 1.012 | 0.909–1.127 | 0.83 |
| S LDL phospholipids, ratio to total lipids | % | 31.58 | 4.10 | 0.990 | 0.887–1.105 | 0.86 |
| XL HDL phospholipids, ratio to total lipids | % | 47.28 | 9.15 | 0.843 | 0.782–0.910 | 1E-5 |
| L HDL phospholipids, ratio to total lipids | % | 48.29 | 4.46 | 1.169 | 1.061–1.288 | 0.002 |
| M HDL phospholipids, ratio to total lipids | % | 47.35 | 3.25 | 1.172 | 1.060–1.296 | 0.002 |
| S HDL phospholipids, ratio to total lipids | % | 54.43 | 4.35 | 0.980 | 0.886–1.084 | 0.69 |

Mean concentrations of the metabolic measures and standard deviations (SD) were averaged across the four cohorts. All 229 lipoprotein, lipid and metabolite measures were quantified using the Nightingale NMR metabolomics platform (Nightingale Health Ltd, Helsinki, Finland). The 14 lipoprotein subclass sizes were defined as follows: extremely large VLDL with particle diameters from 75 nm upwards and a possible contribution of chylomicrons, five VLDL subclasses (average particle diameters of 64.0 nm, 53.6 nm, 44.5 nm, 36.8 nm, and 31.3 nm), IDL (28.6 nm), three LDL subclasses (25.5 nm, 23.0 nm, and 18.7 nm), and four HDL subclasses (14.3 nm, 12.1 nm, 10.9 nm, and 8.7 nm). The mean size for VLDL, LDL and HDL particles was calculated by weighting the corresponding subclass diameters with their particle concentrations. The lipoprotein composition measures were calculated as the ratio of a given lipid species (e.g. triglycerides) to the total lipid concentration within that particular lipoprotein subclass size.

**Supplemental Table 2: Tabulation of all biomarker results in the figures (online spreadsheet).**

**Supplementary Table 3. Regression models for risk of future type 2 diabetes.**

| | Basic clinical score | | | Extended clinical score | | | Biomarker enhanced score | | |
|---|---|---|---|---|---|---|---|---|---|
| **Variable** | **beta** | **SE** | **P** | **beta** | **SE** | **P** | **beta** | **SE** | **P** |
| Male sex | 0.081 | 0.167 | 0.63 | -0.091 | 0.184 | 0.62 | -0.364 | 0.200 | 0.06 |
| Age [years] | 0.028 | 0.015 | 0.06 | 0.033 | 0.015 | 0.03 | 0.0286 | 0.0158 | 0.07 |
| Body mass index [SD] | 0.864 | 0.059 | 8E-48 | 0.681 | 0.072 | 3E-21 | 0.462 | 0.0740 | 4E-10 |
| Fasting glucose [SD] | 0.448 | 0.071 | 3E-10 | 0.343 | 0.068 | 4E-7 | 0.640 | 0.0719 | 5E-19 |
| HDL cholesterol | - | - | - | -0.339 | 0.121 | 0.005 | - | - | - |
| Triglycerides | - | - | - | -0.003 | 0.066 | 0.96 | - | - | - |
| log(fasting insulin) | - | - | - | 0.323 | 0.087 | 0.0002 | - | - | - |
| Large HDL free cholesterol | - | - | - | - | - | - | -0.474 | 0.117 | 5E-5 |
| Phenylalanine | - | - | - | - | - | - | 0.320 | 0.0802 | 6E-5 |
| Large VLDL cholesterol ester % | - | - | - | - | - | - | -0.321 | 0.0847 | 2E-4 |

The weights (beta-coefficients in the multivariable logistic regression models) of three different risk scores for type 2 diabetes were derived by multivariable logistic regression using meta-analysis of three cohorts (YFS, FINRISK-1997, and DILGOM). The 'basic clinical score' was comprised of sex, baseline age, BMI and fasting glucose. An 'extended clinical score' was also examined, which included HDL cholesterol, triglycerides, and fasting insulin in addition to the variables in the 'basic clinical score'. These two models were compared to a 'biomarker enhanced score' comprised of the variables in the 'basic clinical score' and additionally three variables metabolic biomarkers that entered the prediction model (phenylalanine, free cholesterol in large HDL, and cholesteryl esters to total lipids ratio within large VLDL). These three metabolic biomarkers were selected among all clinical risk factors and the NMR-based metabolic measures using a forward step-wise process, which was meta-analyzed across the three derivation cohorts.

**Supplementary Table 4. Risk discrimination of three prediction models for risk of type 2 diabetes, assessed among 5,271 individuals aged 31.**

| Model | C-statistic (95% CI) | Integrated discrimination improvement (IDI) | Continuous net reclassification improvement (NRI) |
|---|---|---|---|
| #1: Basic clinical score | 0.729 (0.692-0.766) | - | - |
| #2: Extended clinical score | 0.752 (0.717-0.786) | - | - |
| #3: Biomarker enhanced score | 0.764 (0.731-0.798);<br>P=0.0003 for #3 *vs* #1<br>P=0.13 for #3 *vs* #2 | #3 *vs* 1:<br>1.32% net (P=7E-9)<br>#3 *vs* # 2:<br>0.51% net (P=0.03) | #3 *vs* 1<br>48.1% net (P=3E-11)<br>#3 *vs* 2<br>22.3% net (P=0.002) |

The absolute risk estimates for assessing discrimination were derived by fitting the weighted sum of risk factors and metabolic biomarkers for each model ("the linear predictor") to the validation cohort (NFBC), and then calculated as

```
Absolute risk = 1/(1+exp(-(slope + scale(linear_predictor)))).
```

The slope and scaled beta-coefficient of the linear prediction was (-3.449; 0.704) for model #1, (-3.546;0.732) for model #2, and (-3.630;0.790) for model #3. The model weights from multivariable logistic as derived in meta-analysis of the 3 derivation cohorts are given in Supplementary Table 3. The DeLong method was used for calculating 95% CIs and P-value comparison of two C-statistics curves. IDI and NRI were calculated as previously described.[9,10] The shuffling of absolute risk separately for individuals who did and did not develop type 2 diabetes during follow-up are illustrated in Supplementary Figure 7.

**Supplementary Figure 1. Overview of study cohorts and participant inclusion.**

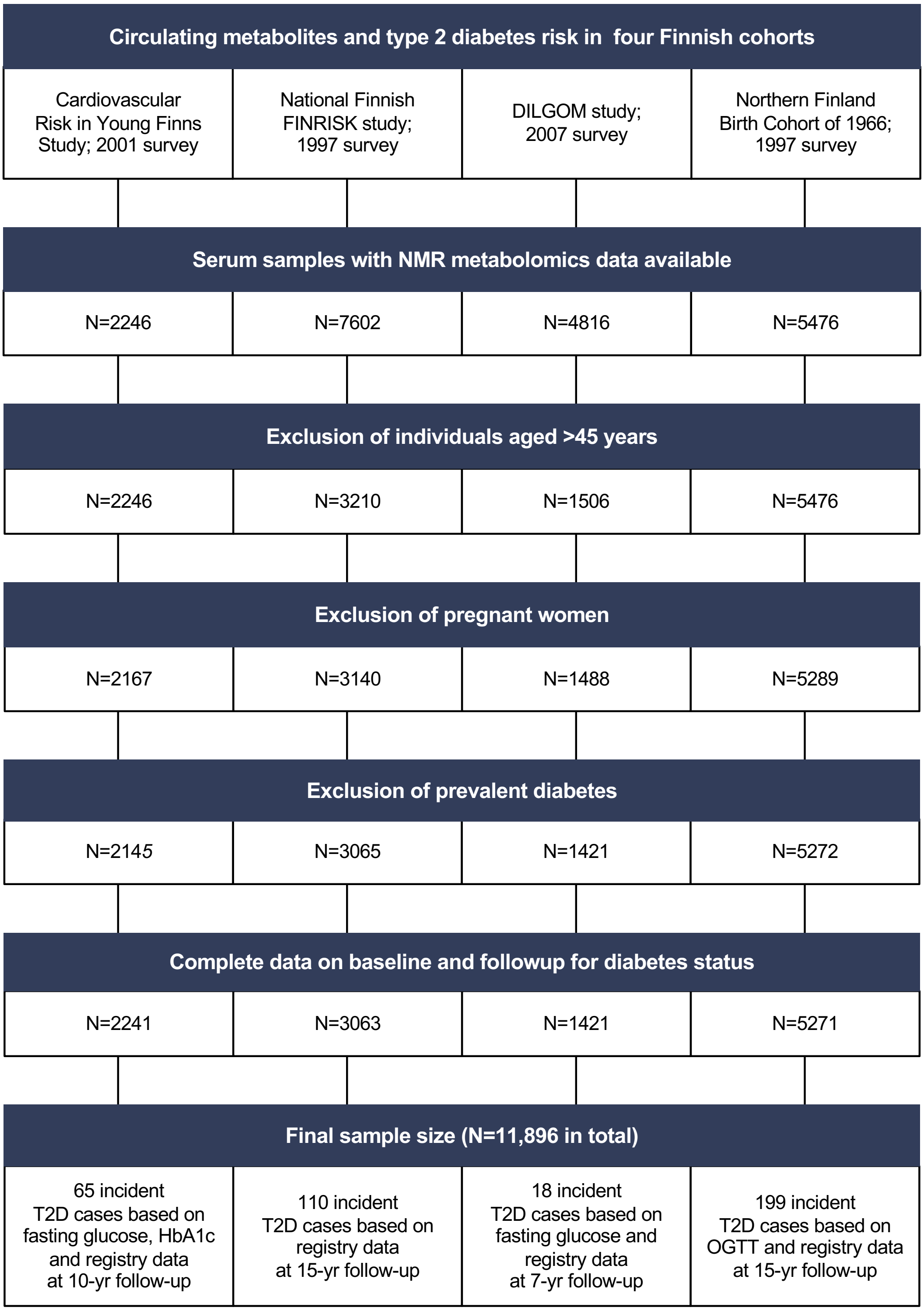


The flow-chart indicates numbers of participants in each cohort after specified exclusion criteria and number of cases of incident type 2 diabetes during 8–15 years of follow-up.

**Supplementary Figure 2. Relation of 125 metabolic measures (not shown in main paper) to risk of future type 2 diabetes.**

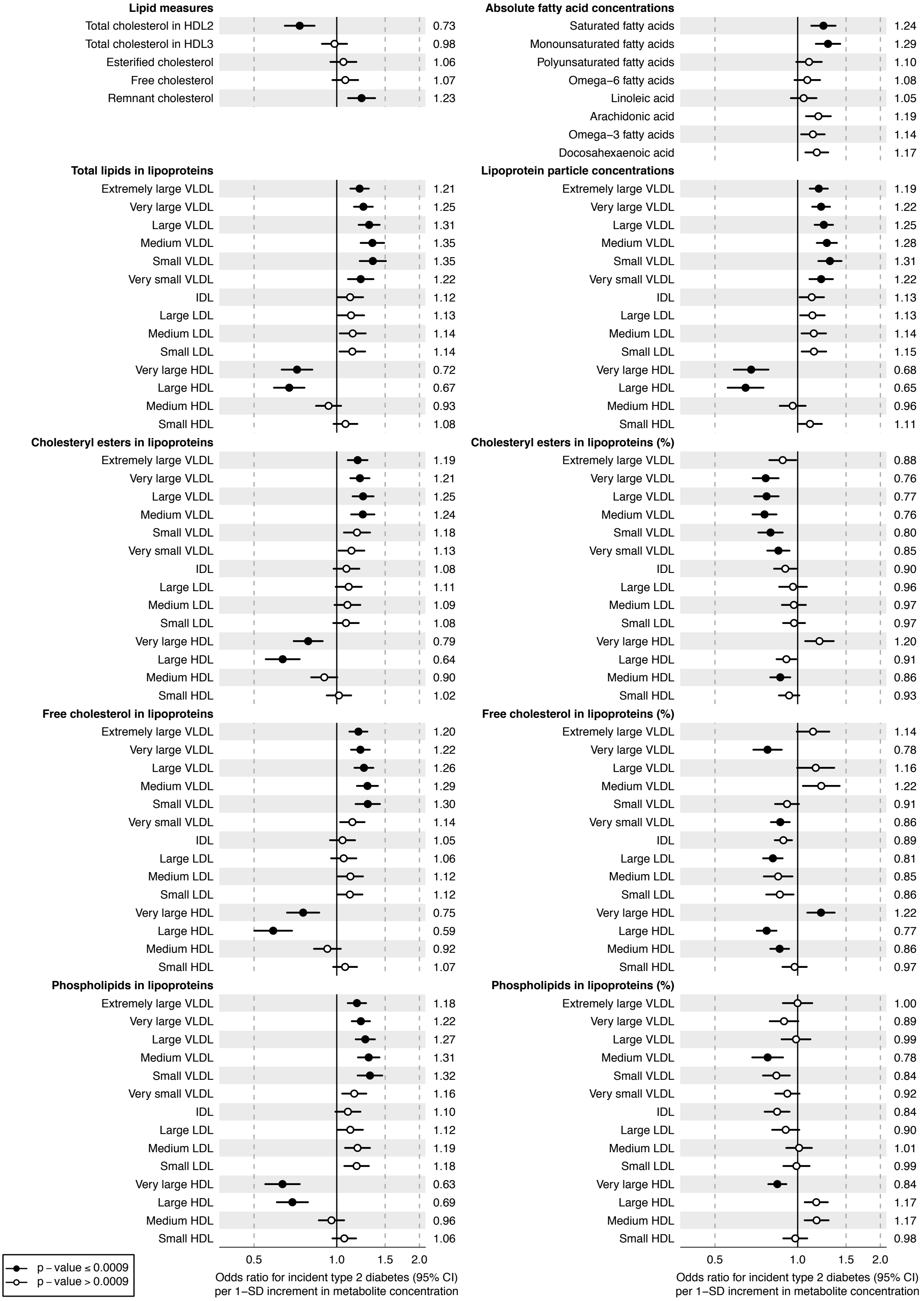


Values are odds ratios (95% confidence intervals) per 1-SD log-transformed metabolite concentration. Odds ratios were adjusted for sex, baseline age, BMI, and fasting glucose. The results were meta-analyzed for 11,896 young adults from four prospective cohorts.

**Supplementary Figure 3. Consistency of biomarkers to the risk of future diabetes across the four cohorts.**

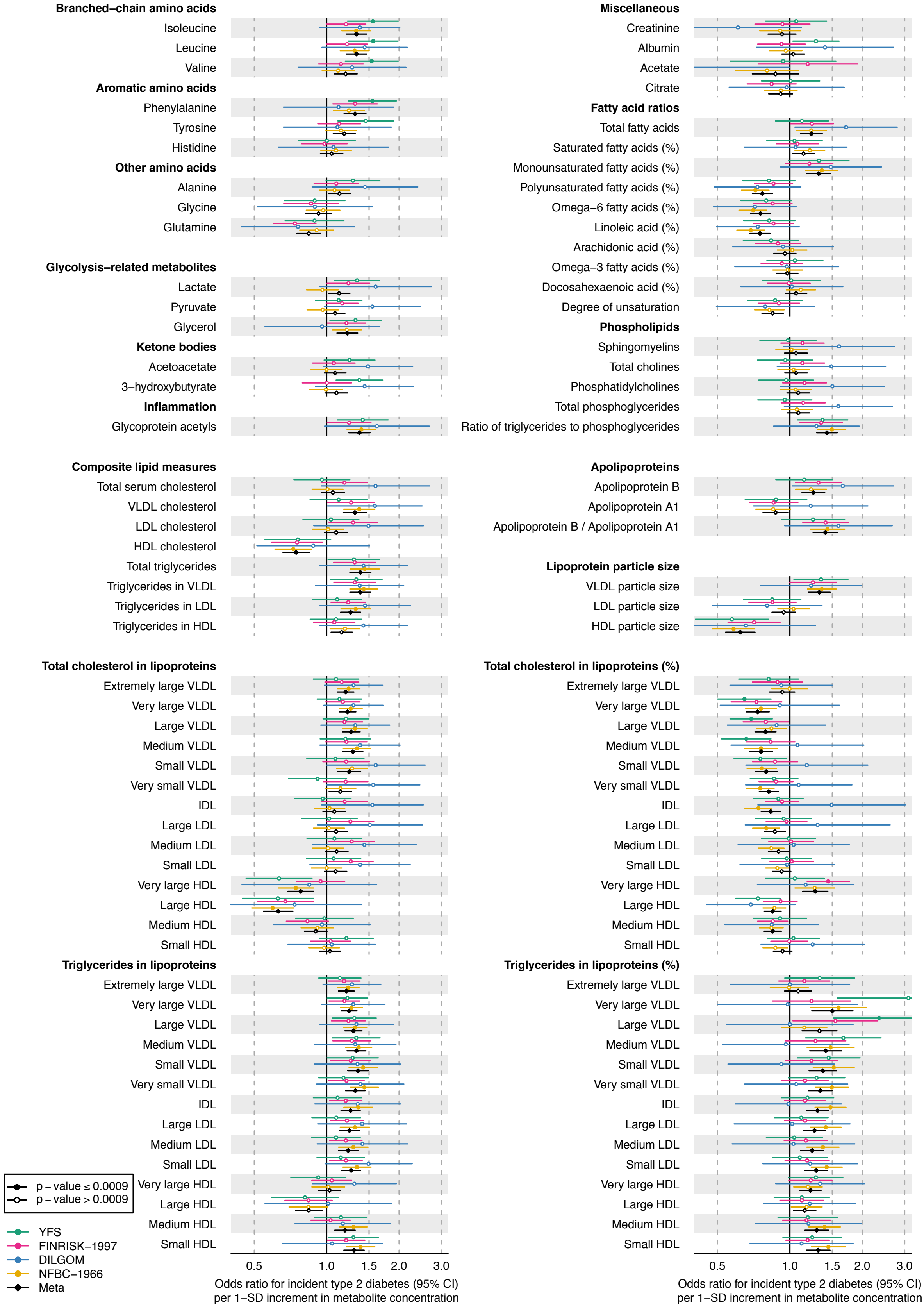


Values are odds ratios (95% confidence intervals) per 1-SD log-transformed metabolite concentration. Odds ratios were adjusted for sex, baseline age, BMI, and fasting glucose. YFS, Cardiovascular risk in Young Finns Study; NFBC, Northern Finland Birth Cohort.

**Supplementary Figure 4. Biomarkers for the risk of future diabetes assessed separately for men and women.**

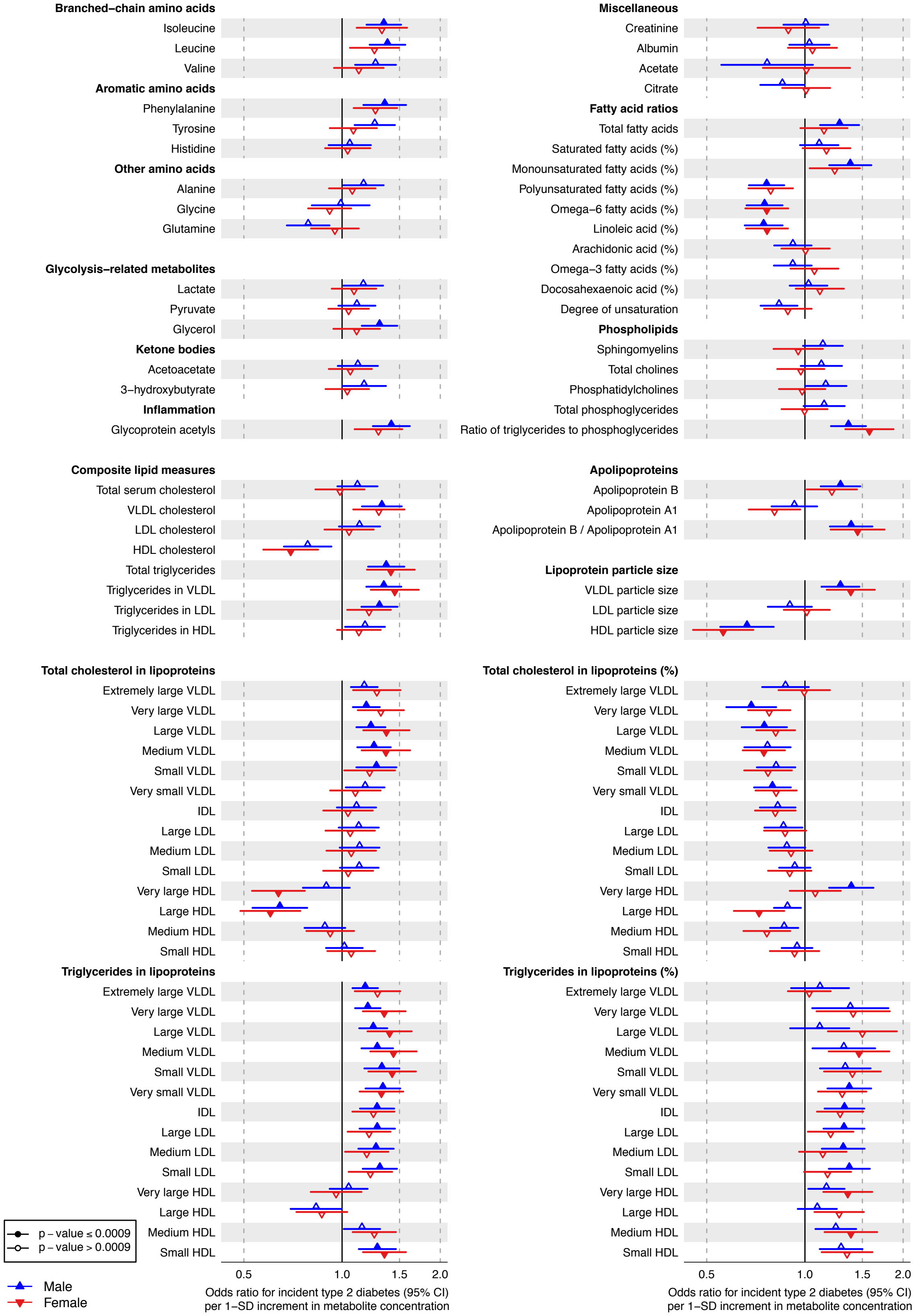


Values are odds ratios (95% confidence intervals) per 1-SD log-transformed metabolite concentration. Odds ratios were adjusted for baseline age, BMI, and fasting glucose. Results were meta-analyzed across the 4 cohorts (5,696 men; 6,200 women).

**Supplementary Figure 5. Biomarkers for the risk of future diabetes compared to cross-sectional associations with BMI, HOMA-IR and fasting glucose.**

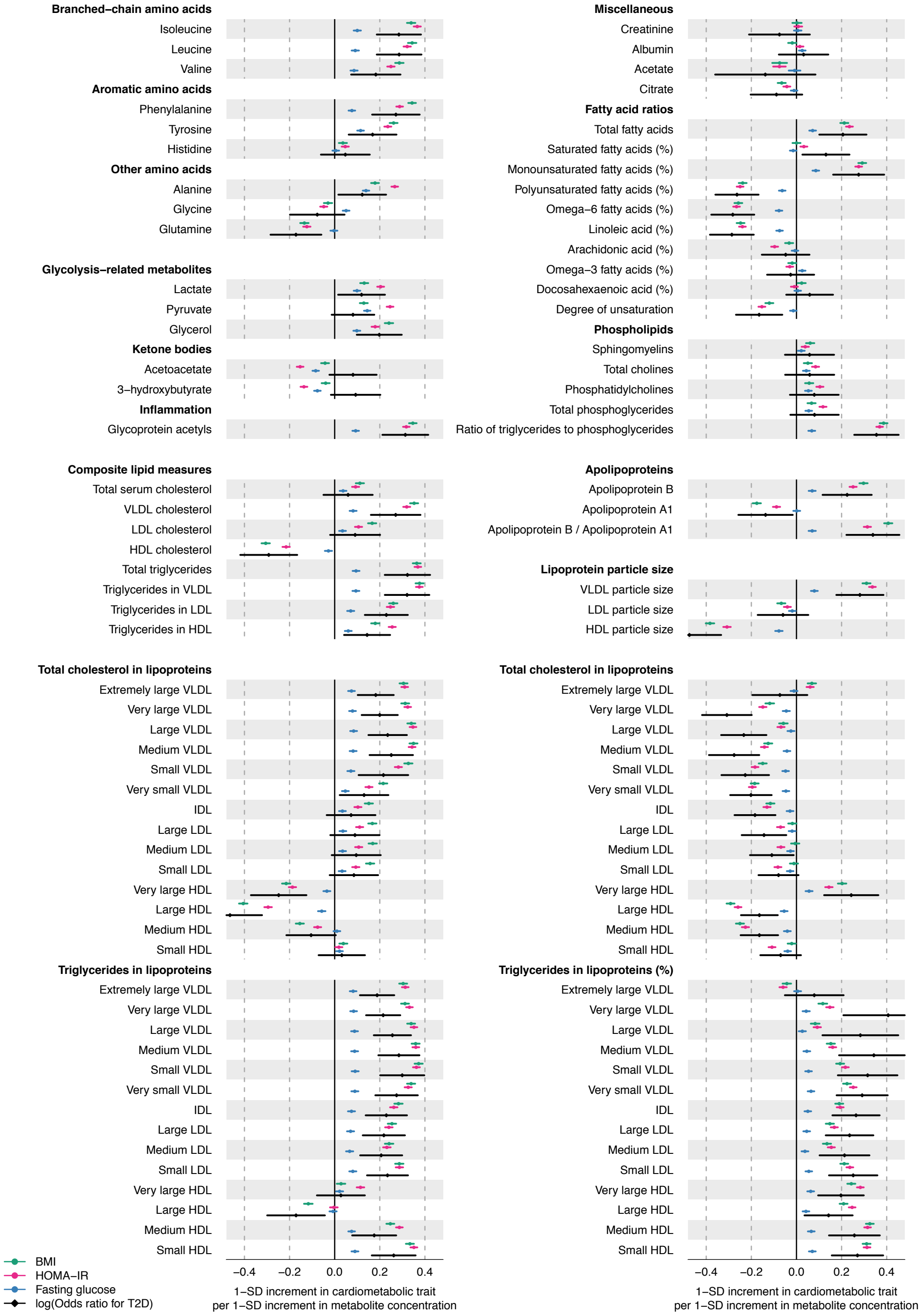


Values are beta-correlations from cross-sectional metabolite associations with BMI (green), log(HOMA-IR) (red) and fasting glucose (blue). To enable comparison of the patterns of associations, magnitudes are scaled to 1-SD in each of the outcomes (corresponding to 4.2 kg/m$^2$ for BMI, 0.57 for log(HOMA-IR) and 0.56 mmol/l for glucose) per 1-SD log-transformed metabolite concentration. Also shown for comparison are the beta-coefficients of the main results for risk of future type 2 diabetes risk (=natural logarithm of odds ratio; black). Results were adjusted for sex and age, and meta-analyzed for 11,896 individuals from the four cohorts. Error bars denote 95% confidence intervals; the large sample size and consistency across cohorts make confidence intervals narrow for the cross-sectional linear regression analyses.

**Supplementary Figure 6. Biomarker associations with diabetes risk after additional adjustment for insulin and without adjustment for BMI.**

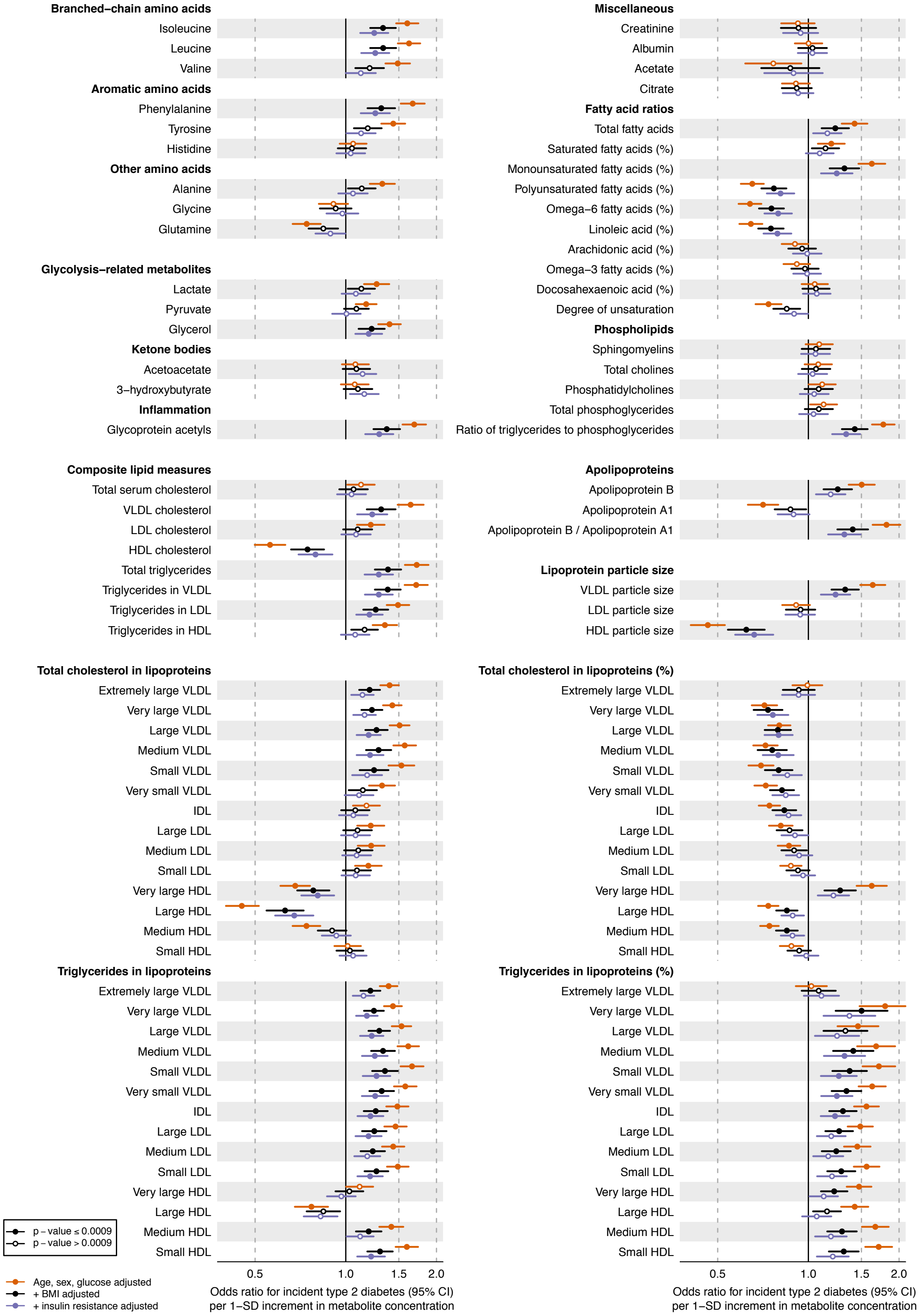


Values are odds ratios (95% confidence intervals) per 1-SD log-transformed metabolite concentration. Odds ratios are shown without adjustment for baseline BMI and for a model adjusting for both BMI and fasting insulin. All models were adjusted for sex, baseline age, BMI, and fasting glucose. The results were meta-analyzed for 11,896 young adults from the four cohorts.

**Supplementary Figure 7. Refinement of absolute risk for type 2 diabetes by including biomarker summary score into prediction model.**

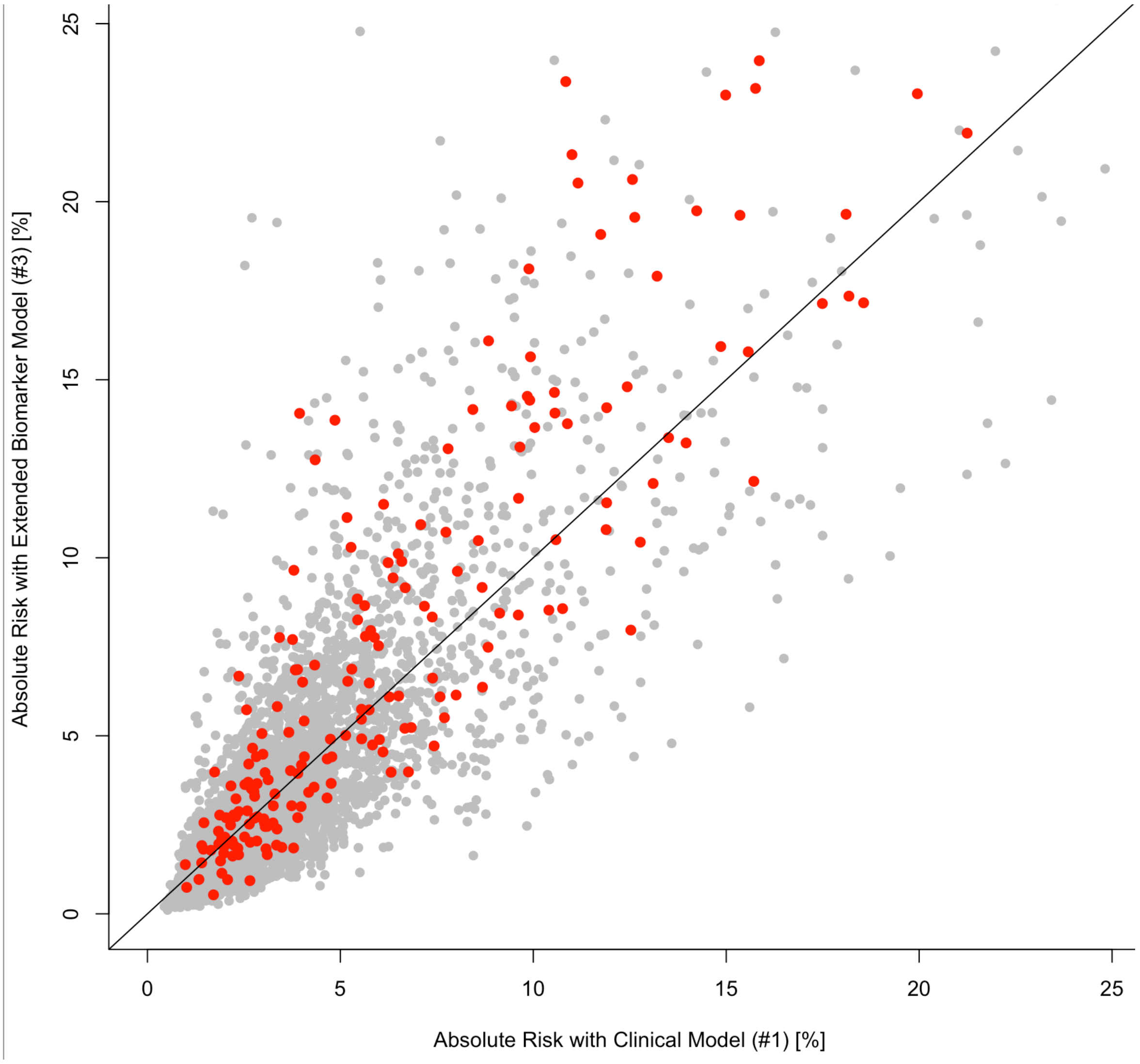


Illustration of the prediction of absolute risk for type 2 diabetes using the 'basic clinical model' compared to the 'biomarker-enhanced model'. The results were assessed in the NFBC validation cohort of 5,271 individuals, who were all 31 years of age at blood sampling. Red dots indicate individuals who developed type 2 diabetes during the 15-year follow-up period, and grey dots indicate those who did not develop type 2 diabetes.
